# Early origin and deep conservation of enhancers in animals

**DOI:** 10.1101/633651

**Authors:** Emily S Wong, Siew Zhuan Tan, Victoria Garside, Gilles Vanwalleghem, Federico Gaiti, Ethan Scott, Edwina McGlinn, Mathias Francois, Bernard M Degnan

**Affiliations:** School of Biological Sciences, University of Queensland, St Lucia, Australia; Institute for Molecular Biosciences, University of Queensland, St Lucia, Australia; Australian Regenerative Medicine Institute, Monash University, Victoria, Australia; School of Biomedical Sciences, University of Queensland, St Lucia, Australia

## Abstract

Transcription factors (TFs) bind DNA enhancer sequences to regulate gene transcription in animals. Unlike TFs, the evolution of enhancers has been difficult to trace because of their rapid evolution. Here, we show enhancers from the sponge *Amphimedon queenslandica* can drive cell type-specific reporter gene expression in zebrafish and mouse, despite sponge and vertebrate lineages diverging over 700 million years ago. Although sponge enhancers, which are present in both highly conserved syntenic gene regions (*Islet–Scaper, Ccne1–Uri* and *Tdrd3–Diaph3*) and sponge-specific intergenic regions, have no significant sequence identity with vertebrate genomic sequences, the type and frequency of TF binding motifs in the sponge enhancer allow for the identification of homologous enhancers in bilaterians. *Islet* enhancers identified in human and mouse *Scaper* genes drive zebrafish reporter expression patterns that are almost identical to the sponge *Islet* enhancer. The existence of homologous enhancers in these disparate metazoans suggests animal development is controlled by TF-enhancer DNA interactions that were present in the first multicellular animals.

**One-sentence summary:** Enhancer activity is conserved across 700 million years of trans-phyletic divergence.

## Background

Metazoan complexity and cell type diversity is contingent upon genome-encoded regulatory information. This information directs cell lineage-specific gene regulatory networks (GRNs) in time and space. The activation and output of GRNs is underpinned by *cis*-regulatory elements, namely promoters and enhancers, which contain short (<10 bp) specific DNA motifs to which transcription factors (TFs) bind. Promoters and proximal regulatory elements originated before the emergence of the animal kingdom, distal regulatory elements appear to be unique to animals (*1*–*3*) (**FIGS1**).

Although the gene expression profiles of tissues and cell types shared across species are highly conserved (*4*), the regulatory elements controlling gene expression appear to be largely species-specific (*5*–*8*). Comparison of the *in vivo* rate of TF binding site evolution with the rate of gene expression change reveals that binding sites evolve at a much faster rate than the TFs they interact with and the expression of the genes they regulate (*9*). TF–DNA binding events are significantly divergent even between human and mouse and a major source of *in vivo* binding differences between human individuals (*5, 10*). For example, only ~5% of profiled *in vivo* embryonic stem cell TF binding sites are conserved between human and mouse (*11*) and only 1% of human tissue-specific enhancers are conserved by alignment of histone marks across mammalian genomes (*6*). Between eumetazoan phyla (bilaterians and cnidarians), only a single conserved enhancer has been uncovered (*12*). Thus, no pan-metazoan enhancer has been described to date despite the fact that the major families of TFs are highly conserved across the animal kingdom.

The widespread conservation of gene order (conserved synteny) serves as indirect evidence of constraints imposed by regulatory elements (*13–16*). The concept of a *cis*-regulatory block was formulated based on the observation that many key developmental genes are enriched at regions of conserved synteny in the human genome (*17, 18*). Almost 600 examples of gene pair conservation (‘microsynteny’) have been identified across metazoans, with 80 reported in the sponge *Amphimedon* genome (*16*). Genes within conserved regulatory blocks are classified into one of two types, either ‘bystander’ – the genes assumed to harbor regulatory sequences, or developmental – the candidate targets of *cis*-regulatory elements (**FIG1A**). The hypothesized model for this strict maintenance of gene arrangement across evolution is an interleaving regulatory structure, whereby the regulatory elements within the span of one gene serves as input to an adjacent gene, which results in the misregulation of the target gene if the bystander gene is uncoupled or interrupted (*19*) (**FIG1A**). The regulatory enhancers purported to underpin the conservation of these microsyntenic units across the animal kingdom have yet to be characterized, largely because of the rapid evolution of these non-coding sequences (**FIGS1C)**.

**Figure 1.**
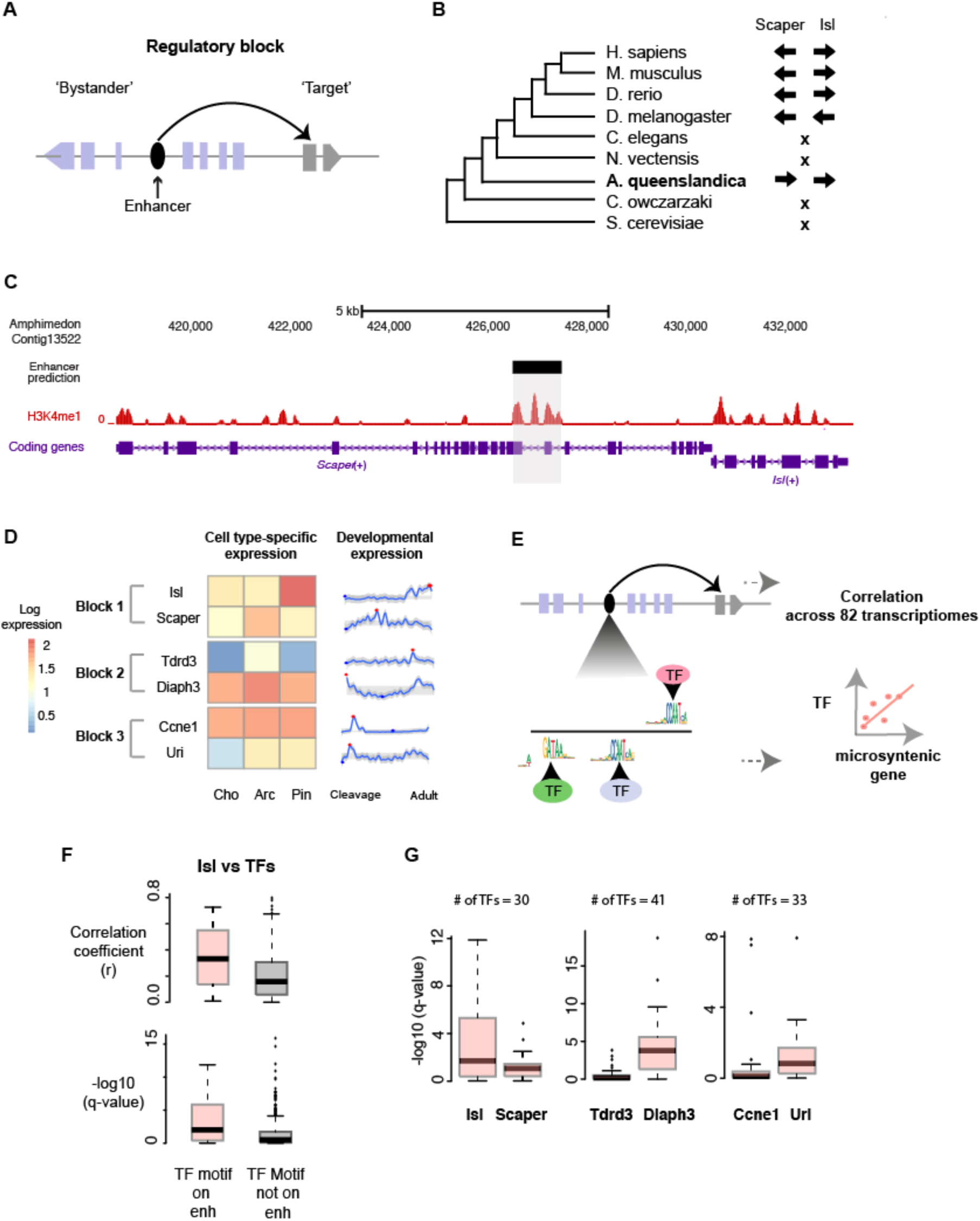
Conserved microsyntenic regions in sponge and vertebrates have similar regulatory features. **(A)** Structure of a genomic regulatory block. Conserved microsyntenic regions are thought to be maintained because regulatory elements residing in one gene serve as regulatory input to an adjacent ‘target’ gene, commonly a developmental TF (*19*) **(B)** The genomic arrangement of *Islet* and *Scaper* is conserved in animal genomes, including in *A. queenslandica*. Arrows, direction of transcription; **x**, *Islet*-*Scaper* synteny not present. **(C)** The location of a putative enhancer in *Amphimedon Scaper* (grey) overlaps with a H3K4me1 peak. **(D)** The expression levels of genes comprising three microsyntenic block in adult *Amphimedon* cell types (Cho=choanocytes, Arc=archeocytes, Pin=pinacocytes) and during development from early embryo to adult. Expression levels are log_e_ transformed counts. Red and blue dots in the timecourse refer to the maximum and minimum values, respectively. Grey region in time course denotes the interquantile range. **(E)** Prediction of TF binding motifs in an enhancer by position-weighted matrices (PWMs) and using gene expression data to test if the predicted set of TFs and the candidate target gene share gene regulatory networks. **(F)** Developmental gene expression are significantly higher correlated between *Amphimedon Islet* and the TFs whose binding motifs are present in the *eISLs* enhancer (n=30) compared to other TFs (n=454) (Mann-Whitney U, p=5.5e-4). Pearson’s correlations were adjusted for multiple testing (‘q-value’). **(G)** Predicted sponge TFs (number at the top of each plot) associated to the *Islet-Scaper* regulatory unit were better correlated with developmental gene *Islet* than *Scaper* (Mann-Whitney U, p<0.05). TFs associated with the enhancer in *Diaph3* showed higher correlations with *Diaph3* than *Tdrd3*. TFs associated to the enhancer in *Ccne1* were better correlated for *Uri* than *Ccne1*.

Although enhancer sequences evolve rapidly, they can operate with substantial sequence flexibility suggesting that regulatory conservation may be more common than realized. Numerous studies have shown that strict and specific TF binding motif positional conservation is not a requirement for enhancer activity (*20*–*22*). Between distant fly species, *eve* stripe 2 enhancer sequences show extensive flexibility in TF binding motif arrangement and spacing, and yet show the same expression activity pattern (*20*). Similarly, extreme sequence divergence has been reported in orthologous regulatory regions that produce similar regulatory output in yeast (*23*), and individual TF binding motifs in yeast and mouse have been show to tolerate up to 72% of all possible mutations (*24*).

Sponges are considered one of the oldest surviving phyletic lineages of animals diverging from other metazoans around 700 MYA. To understand the origin and evolution of animal *cis*-regulatory elements, we analysed the activity of enhancers from the sponge *Amphimedon queenslandica*, which are located within and outside microsyntenic regions, in zebrafish and mouse transgenic reporter lines. These sponge enhancers are able to drive cell-type specific reporter gene expression in these vertebrates. From the analysis of the type and frequency of TF binding motifs in sponge enhancers we are able to identify homologous enhancers in humans and mice. These human and mouse orthologous enhancers drove highly similar zebrafish expression patterns to that of the sponge enhancer. These findings reveal an unexpectedly deep conservation of metazoan enhancers despite high sequence divergence.

## RESULTS

### Characterization of sponge enhancers within conserved syntenic regions

*Amphimedon* enhancers were identified by genome-wide chromatin immunoprecipitation of antibodies for the histone modifications H3K4me3, H3k27ac, H3K4me1, H3K36me3 and H3k27me3 followed by high-throughput sequencing (ChIP-Seq) and alignment of the short-reads to the *Amphimedon* genome (*2, 13*). Enhancers within microsyntenic regions were found by overlaying the locations of metazoan syntenic units with enhancer locations inferred from a classification of the histone marks (*16*) (e.g. **FIG1B**). 43 of the 80 reported metazoan syntenic units were identified, where least one sponge candidate enhancer was active during larval or adult stages (*2*). These putative sponge enhancers do not have significant sequence identity to any regions in known vertebrate genomes. Of these 43, we have focused on three enhancers located within the conserved syntenic units of the gene pairs: *Islet–Scaper, Tdrd3–Diaph3* and *Ccne1–C19orf2/Uri* (**FIG1C, FIGS2**). These candidate enhancers were selected based on unambiguous phylogeny, strength of H3K4me1 enhancer signal, and known developmental regulatory roles for the linked genes in these microsyntenic regions.

To infer whether sponge genes within these regulatory units show evidence of discrete functional roles, we compared gene expression for syntenic gene pairs across development, under the expectation that microsyntenic genes with different functions will display decoupled patterns of expression. We determined the gene expression level of *Amphimedon* genes spatially and temporally by performing single cell RNA-Seq (CEL-Seq) on three adult sponge cell types (pinacocyte, archeocyte and choanocyte) and by constructing a dense 82 stage developmental time series from early embryo, to swimming larva, to sessile adult (*25*)(**FIGS3A**). We found clear differences in spatial and/or temporal expression profiles between all three target and bystander gene pairs (**FIG1D**), consistent with reports of uncorrelated expression between target and bystander genes in human and zebrafish (*16, 19*).

Although sponges lack the diversity of cell types present in bilaterians (e.g. neurons, muscles), the sponge genome is surprisingly complex and contains almost all bilaterian TF families (*13, 26*)(**FIGS1B, S3B**). Based on the notion that a gene regulated by the enhancer may show covariance in gene expression with the TFs regulating it, we sought to determine whether TF binding motifs could predict the repertoire of TFs recruited to an enhancer (**FIG1E-G, S4)**. We used position weighted matrix (PWM), an established method to identify TF binding sites (*27, 28*). 117, 160 and 131 motifs were identified for enhancers found in *Islet–Scaper* (*‘eISLs’*), *Tdrd3– Diaph3* (‘*eTDRs’*) and *Ccne1–Uri* (‘*eCCNs’*), respectively at a p-value<1e-3 (**Methods, SM**). Where possible, these motif PWMs were associated to sponge TFs (**Methods, SM**). Within each microsyntenic unit, the set of TFs identified by their binding motifs in the enhancer sequence was significantly more predictive of the gene expression profile of one gene in the microsyntenic block compared to the other (**FIG1G, S4**). For example, TFs associated with the enhancer located within the *Scaper* gene were more strongly correlated with the *Islet* expression profile than *Scaper* profile (Mann-Whitney U, p<0.05). Further, correlations with microsytenic genes were most different when the matched regulatory enhancer was used for prediction consistent with different usage of the same element for alternative functions (*29*). These results demonstrate that PWMs can meaningfully identify those TFs that likely regulate gene expression in *Amphimedon* without prior experimental evidence of the TFs recruited to the enhancer.

### Sponge enhancers direct cell tissue-specific expression in vertebrates

We assessed the activity of the three sponge enhancers at metazoan-conserved syntenic regions in transgenic zebrafish. Enhancer reporter assays in zebrafish are routinely used to detect enhancer activity *in vivo* (*30, 31*), and reporter expression reflects that of the endogenous gene (*32*). We inserted the sponge enhancer sequence upstream of a silent *gata2a* promoter and GFP reporter gene within a Zebrafish Enhancer Detection (ZED) vector (*33*) (**Methods)**. Independent stable transgenic lines were generated for each construct (**FIG2A, TableS1-2**).

**Figure 2.**
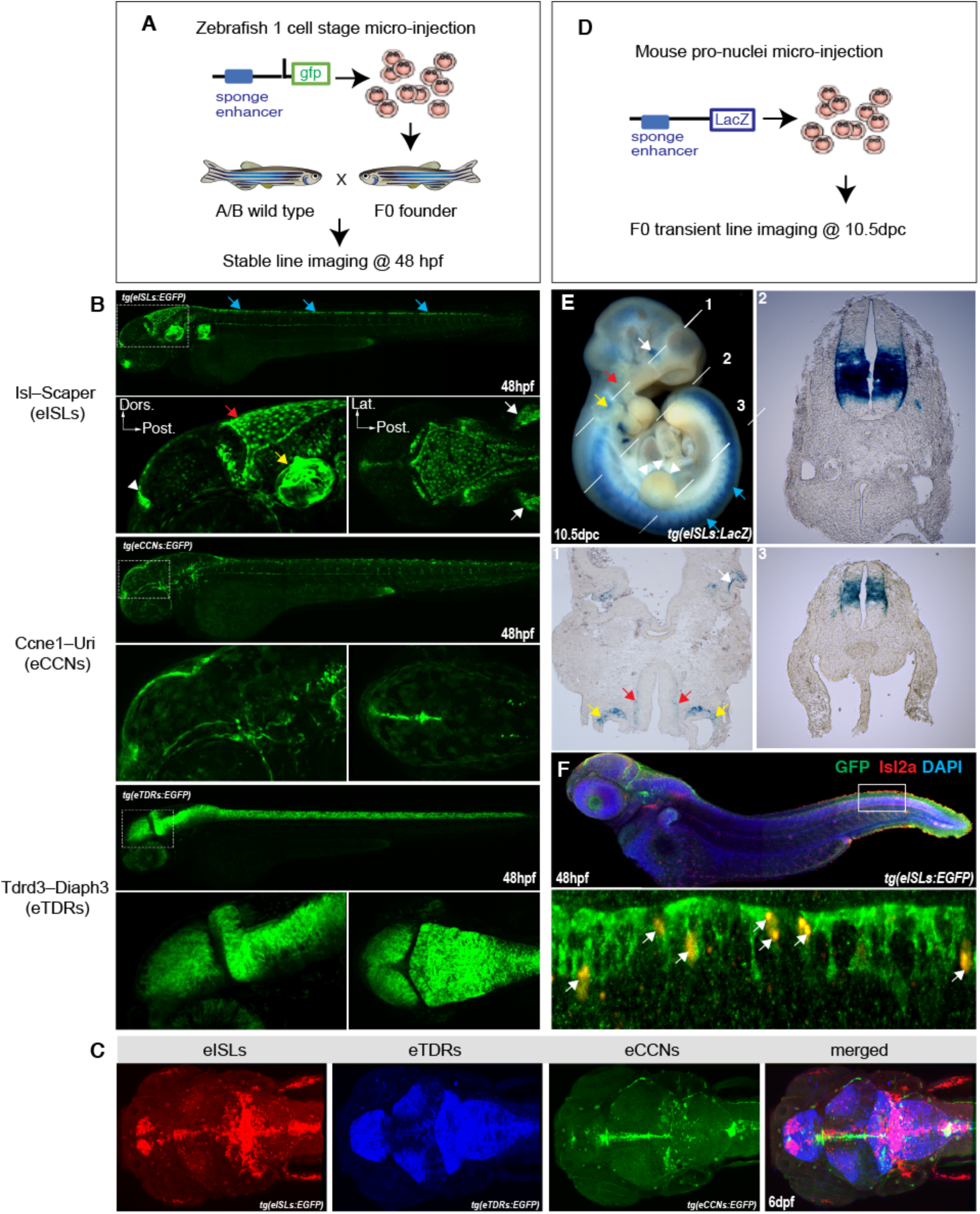
Sponge enhancers direct cell-type specific expression in vertebrate embryos. (**A**) Schematic of the transgenesis approach used in zebrafish model system. Sponge enhancers are driving a GFP reporter gene in an enhancer trap assay. Three stable transgenic lines for each sponge enhancers from conserved syntenic regions were established: *Isl2–Scaper* (*eISLs*), *Ccne1–Uri* (*eCCNs*) and *Tdrd3–Diaph3* (*eTDRs*). (**B**) Expression pattern of enhancer-driven GFP during zebrafish larvae development at 48hpf: eISLs activity is detected in neuro-epithelial layer of the hindbrain (red arrows), the roof plate (blue arrows), the otic vesicle (yellow arrows), the pineal region (arrowhead) and the pectoral fin (white arrows). eCCNs shows similar activity but in not in identical regions than eISLs with some additional signal in the peri-orbital area. eTDRs reveals pan-neural activity in the spinal cord and in the midbrain. (**C**) Neuroanatomical visualization of the three sponge enhancer activities by image alignment to a common template built from the confocal images (‘merged’). There is a small subset of neurons in the cerebellum in which all three sponge enhancers are active (white signal in ‘merged’). (**D**) Schematic of the mouse transgenesis approach used to generate F0 transient lacZ reporter embryos. **(E)** The *eISLs* sponge enhancer is driving a beta-galactosidase reporter construct. Transient transgenic embryos were harvested at 10.5dpc and whole-mount beta-gal staining revealed LacZ activity in the hindbrain (red arrows), apical-ectodermal ridge of the limb bud (arrowheads), otic vesicle (yellow arrow), spinal cord (blue arrows) and the inner layer of the optic cup (white arrow) in 2 out 4 transgenics. Transverse sections (white dashed line 1,2 and 3) show specific activity of eISL enhancer in a subset of neuron in the neural tube along an antero-posterior axis. (**F**) Double immunofluorescence and *in situ* hybridization for GFP (green) and endogenous*Isl2a* (red) in *eISLs* transgenic reporter fish at 48hpf (upper panel) shows a sub-population of neurons from the roof plate expressing both the GFP and endogenous *Isl2a* mRNA (arrows, lower panel). DAPI is shown in blue to highlight overall tissue organization.

All three *Amphimedon* enhancers drove highly specific reporter expression during zebrafish development with all enhancers driving only shared expression in the roof plate neurons. We imaged the fish by confocal microscopy at 24, 48 and 72 hours post-fertilization (hpf). The 709 bp *eISLs* enhancer was active in the neuro-epithelial region in the mid and hindbrain, a subset of sensory neurons in the roof plate around the midline as well as in the pectoral fin and otic vesicle. Reporter expression was detected from 24 hpf and persisted in these regions until at least 72 hpf (**FIG2B-C, S5**). Similarly, the 931 bp *eCCNs* and the 1,510 bp *eTDRs* enhancers appeared active in neuronal lineages (**FIG2B-C**). *eTDRs* showed a pan-neural activity labeling most neurons from the spinal cord and mid brain region while *eCCNs* labeled a small peri-orbital neuron population (**FIG2B-C**).

To firmly establish the tissues associated with enhancer activity, we performed RNA-Seq on GFP-positive and negative cell fractions from our stable fish reporter lines (**Methods, FIGS6**). By mapping functional terms associated with significantly upregulated genes in reporter-positive cells, we showed each sponge enhancer labeled a highly specific population of cells paralleling their observed anatomical location. For example, genes linked to otolith, ear development, notochord, melanocyte, and fin development genes were enriched in *eISLs* positive cells supporting the reporter gene localization in the ear, spinal cord, skin and pectoral fin, respectively (p<0.05, over 5 fold difference) (**Methods**).

Previous work has reported a candidate zebrafish enhancer of *Isl2a* (an ortholog of sponge *Islet*) intronic to *Scaper* (*16*). This putative zebrafish endogenous enhancer was studied by transgenic reporter fish assay, with reporter expression displaying a similar expression pattern to the endogenous *Isl2a* transcript (*16*) (reproduced in this study, **FIGS7**). Hence, we sought to assess whether the sponge *eISL*s enhancer was also active in the same cells and at the same time as the endogenous *Isl2a* transcript. To dissect coexpression at cellular resolution, we performed fluorescent *Isl2a in situ* hybridization combined with GFP immunofluorescence in 48 hpf *eISLs* reporter zebrafish embryos. Endogenous *Isl2a* expression was consistent with published expression patterns (*34*), and importantly, was coincident with GFP reporter in distinct sensory neurons in the roof plate and hindbrain regions (**FIG2F**; white arrows), suggestive of homology between the zebrafish and sponge enhancer. Additionally, numerous cells were found to be GFP-positive without endogenous *Isl2a* expression, suggesting the *eISLs* element may also serve as a regulatory sequence for TFs involved in the transcriptional regulation of genes other than *Isl2a* (**FIG2F)**.

We further sought to determine whether the same sponge enhancers could display similar cell type specific activity in mammals. To this end, we characterized the expression profile of LacZ reporter constructs in transient transgenic mouse embryos at 10.5 days post coitum (dpc) (**FIG2D, Methods**). We found the sponge *cis*-regulatory elements directed beta-galactosidase activity in a tissue-specific manner in mouse. These activity patterns were consistent with the expression patterns observed in fish reporter assays (**FIG2E, FIGS8**). For instance, beta-galactosidase activity for *eISLs* was detected in the mouse neural tube, otic vesicle, limb bud and hindbrain, suggesting that the sponge enhancers are active in largely conserved tissue-specific gene regulatory networks in vertebrates.

Given all the tested sponge microsyntenic-derived enhancers all appear to direct gene expression in vertebrates, we sought to determine if enhancers in non-conserved genomic regions in the sponge also functioned in vertebrates. To achieve this, we selected two novel enhancers located within protein-coding genes that are sponge-specific, lacking 1-to-1 vertebrate orthology (termed ‘*e1s’*, ‘*e2s’*) (**FIGS9**). Interestingly, compared with enhancers in conserved syntenic units, we found these intragenic enhancers consistently drove broader reporter expression in stable fish lines (**FIGS10)** and transient transgenic mouse (data not shown). Of note, no unambiguous GFP signal was detected in animals injected with empty vectors containing the minimal promoter and gfp sequence (**FIGS11**).

In summary, despite a lack of primary sequence similarity, *Amphimedon* enhancers within microsyntenic regions are active in teleost and mammalian development and direct restricted cell-type specific activity. Interestingly, sponge enhancers outside of conserved syntenic regions, and which likely regulate sponge-specific genes, were still able to respond to TF activation in vertebrates, suggesting the combinations of transcription factor binding sites (TFBS) on sponge enhancers can be broadly interpreted across metazoans.

### Identification of mammalian orthologs of a sponge enhancer

Given the ability of sponge enhancers to drive cell type-specific expression in zebrafish despite ~700 million years of independent evolution, we sought to determine if sponge enhancers possess a conserved *cis*-regulatory signature that could be used to identify orthologous enhancers in bilaterians despite the absence of detectable sequence similarity.

Given a difference in the TF binding motif signature of enhancers compared to background (**FIG3B)** and that TF binding motifs at sponge enhancers are able to predict interacting TFs (**FIG1F**), we devised a method to align divergent enhancers based on their collection of binding motifs (**FIG3A**). Using a sliding window approach, we scanned bilaterian microsyntenic regions with sponge enhancer sequences measuring sequence similarity by the type and frequencies of TF binding motifs; motif order and orientation were ignored (**Method**). We assumed that for enhancers to be maintained across large evolutionary distances, their regulatory syntax must allow for extensive flexibility in motif ordering and spacing.

**Figure 3.**
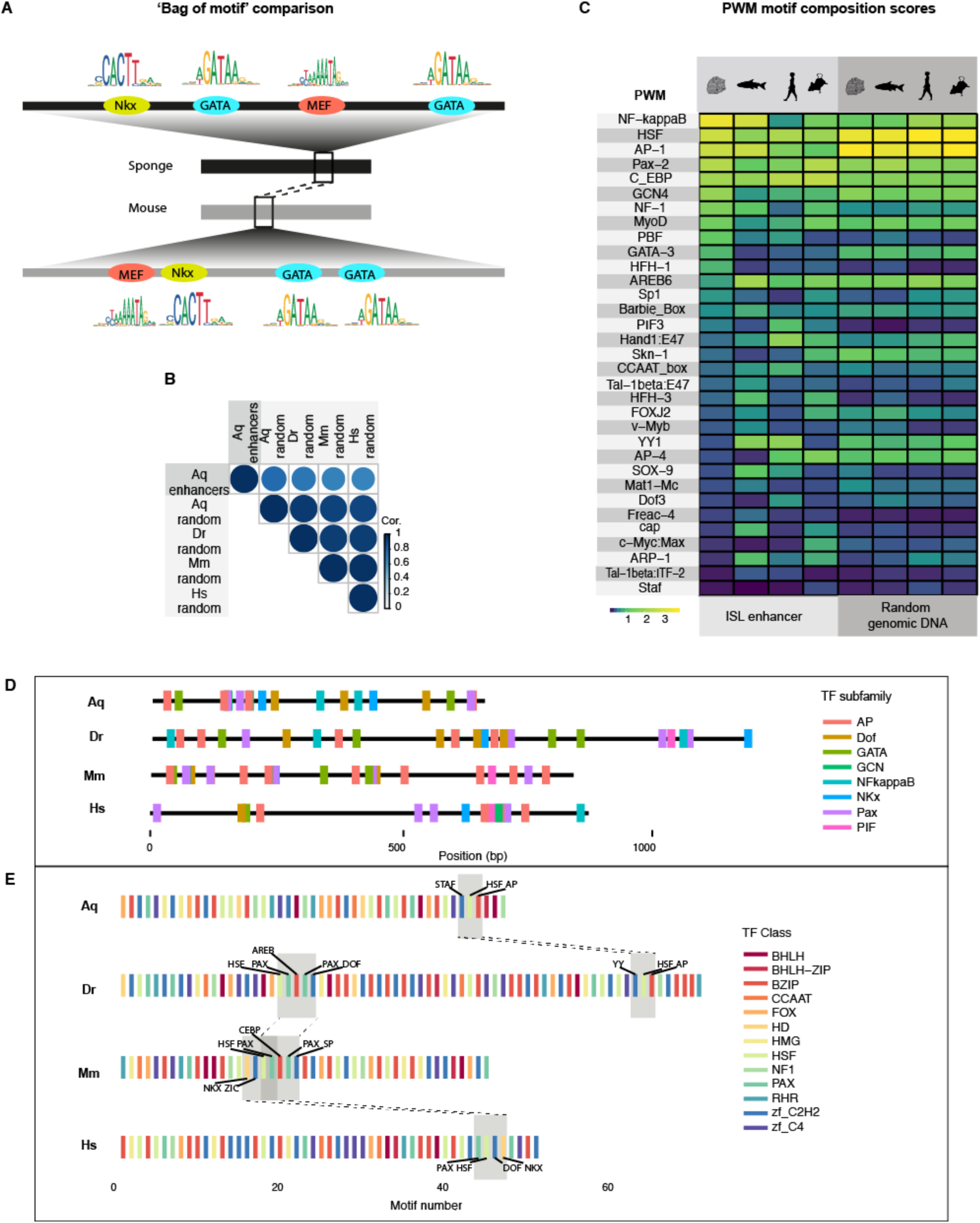
TF motif composition to extract sequence divergent orthologous enhancers. **(A)** Simple representation of the method to compute similarity between divergent enhancer sequences. The metric is based on the type and frequency of TF binding motifs from position weight matrices (PWMs) (**Method**). **(B)** Pearson’s correlation of motif composition scores between *Amphimedon* enhancers by chromatin state (n=200; random subset of length 400bp) and sequences sampled randomly from *Amphimedon* (‘Aq’), zebrafish (‘Dr’), mouse (‘Mm’) and human (‘Hs’) genomes (500 sequences of 1kb from each genome). Scores were L2 normalized to adjust for sequence length difference. Scores quantitate the motif frequency and the statistical significance of the aligned PWM (250 PWMs at motif alignment p<1e-3). **(C)** TF binding motif scores for individual motifs (y-axis) of homologous *Islet* enhancer sequences with 500 randomly sampled sequences of 1kb from matched genomes (mean signals shown). Scores reflect motif frequency and the statistical significance of the aligned PWM. Scores were weighted by PWM significance in the sponge query sequence, under the assumption highly scored binding motifs in sponge are more likely to be functionally relevant. **(D)** Linear representation of 10 most highly scored sponge motifs represented as TF subfamily across homologous *Islet* enhancers. Figures based on alternative filtering of TF motifs in **Fig S12. (E)** PWM motifs (p<1e-3) from putative Islet homologs by TF class (*49*). Grey regions connected by dotted lines are locally aligned p<0.1. TF motifs for these regions are annotated. Only shared motifs amongst sequences are depicted.

First, to identify a human region homologous to *Amphimedon eISLs*, we tested this approach on the human *Scaper*-*Islet* microsyntenic sequence (557 kb). The best-matched region in humans was chr15:76358600-76359500 (900 bp), partially overlapping an enhancer chromatin state in adult lung (*35*). This sequence aligned to the mouse genome at chr9:55560636-55561298 (UCSC Genome Browser) near the 3’ end of the *Scaper* gene in a region of conserved exon synteny to the *Isl2a* candidate enhancer in zebrafish. Based on mouse ENCODE, the region was marked by H3K4me1 (E10.5) in the face and midbrain. It is worth noting only a small fraction of mouse *Scaper*-*Islet* region showed potential enhancer activity in brain, with 11 H3K4me1 peaks in face (~6kb in total; <0.02% of total search region) and 9 in midbrain (~5kb; ~0.01%). The predicted human and mouse homologs of sponge enhancers, *eCCN* and *eTDR*, can be found in **SM**.

We characterized the TF binding motifs composition of the possible *eISL* enhancers in sponge, fish, human and mouse. Comparison of our scores across *Islet* enhancer homologs revealed high variability (**FIG3C, FIGS12**). We accounted for expanded TF families in bilaterians compared to sponge (**FIG3D**), identifying 39 binding motifs common to all species (motif discovery threshold p<1e-3) (**FIGS13A-B, SM)**. Of these, 29 motifs were single TF binding sites across 13 classes of TFs including, for example, leucine zipper factor (bZIP) motifs from AP-1 and C/EBP, and homeodomain family (HD) domain NKx factors. We observed high flexibility in the type and spacing of motifs across all sequences.

The relative position of TF binding sites along an enhancer can be critical as this can change the secondary structure of the DNA and effect the binding of cofactor proteins (*36*–*38*). Thus, we sought to test for the conservation of motif order between these enhancers. We determined the optimal alignment of motifs between the sequences by pairwise sequence alignment and generated empirical null distributions by scrambling the order of motifs in the target (**Method**). We found evidence of motif order conservation (p<0.05) by global-local alignment between the sponge and human enhancers, and between the sponge and mouse elements (p=0.02 and p=0.04, respectively). The aligned regions were short comprising of 2-3 motifs and the best-aligned regions between sponge and human did not match best-aligned regions in the sponge–mouse alignment (**SM**). Hence, we further assumed a more relaxed model based on TF classes to detect conserved motif order. We compared common motifs between the four sequences to 13 protein-binding domains, but did not identify a common local match across all pairwise comparisons (**FIG3E**). Taken together, the results are consistent with a flexible evolutionary model where TF binding sites can be progressively gained or lost while maintaining cell-type specificity in regulatory output.

Given our orthologous predictions have been centered upon chordates, we sought to determine a candidate ortholog in a different phylum. Using *eISLs* as a query sequence in our method described above, we extracted the best aligned fruit fly (*D. melanogaster*) region (800bp) within the genomic span of the fly ortholog to *Scaper* ‘ssp3’ (chr2L:18920300-18921100 (dm6); 35 test regions; **FIGS14, Methods**). Using single cell ATAC-Seq data at 6-8 and 10-12 h after egg laying (*39*), we found the region showed open chromatin in cells of the central nervous system (CNS) – similar tissues to where the fly *Islet* ortholog (‘*tup’*; FlyAtlas) is expressed. The element also showed conserved synteny relative to the 3’ terminal end of fly *Scaper* ortholog (‘*ssp3’*) with vertebrate candidate *Islet* enhancers.

### Predicted mammalian enhancers show highly comparable expression patterns to sponge

We validated the *eISL* enhancer activity of predicted human and mouse sequences in zebrafish and showed both candidate enhancer homologues were able to direct GFP reporter expression in a similar manner to their sponge enhancer counterpart. To assess homology of functional activity between mammal and sponge enhancers, we made stable fish lines for the human and mouse *ISL* enhancers. Cross-species comparisons revealed all enhancers displayed expression pattern in the sensory neurons of the roof plate, as well as in the hindbrain neuro-epithelial layer, the pineal region and the ear capsule (**FIG4A-C, S16**). The sponge *eISL* regulatory elements showed robust activity in neurons of the pectoral fin whereas mouse and human *eISL* orthologs appeared weak in this particular tissue (**FIG4B**; dorsal view pink arrows). Of note, both the mouse and the human *eISL* enhancers displayed specific activity in the eye tissue – mouse *eISL* activity was detected in a polarised population of neurons in the anterior part of the retina (**FIG4A, b**; yellow arrowheads), whereas human *eISL* showed activity in the central part of the lens (**FIG4A, b;** red arrowheads) and in the proctodeum (**FIG4A**; asterisks). Lastly, only the human *eISL* enhancer displayed a non-neural restricted activity and governed GFP expression in a subset of endothelial cells from the inter-somitic vasculature (**FIG4B**; lateral view 2 red arrows). When sponge and mouse enhancer reporter signals were overlaid in a common reference template, this reveals a subset of neurons in which both regulatory regions are active (FIG4c; white signal).

**Figure 4.**
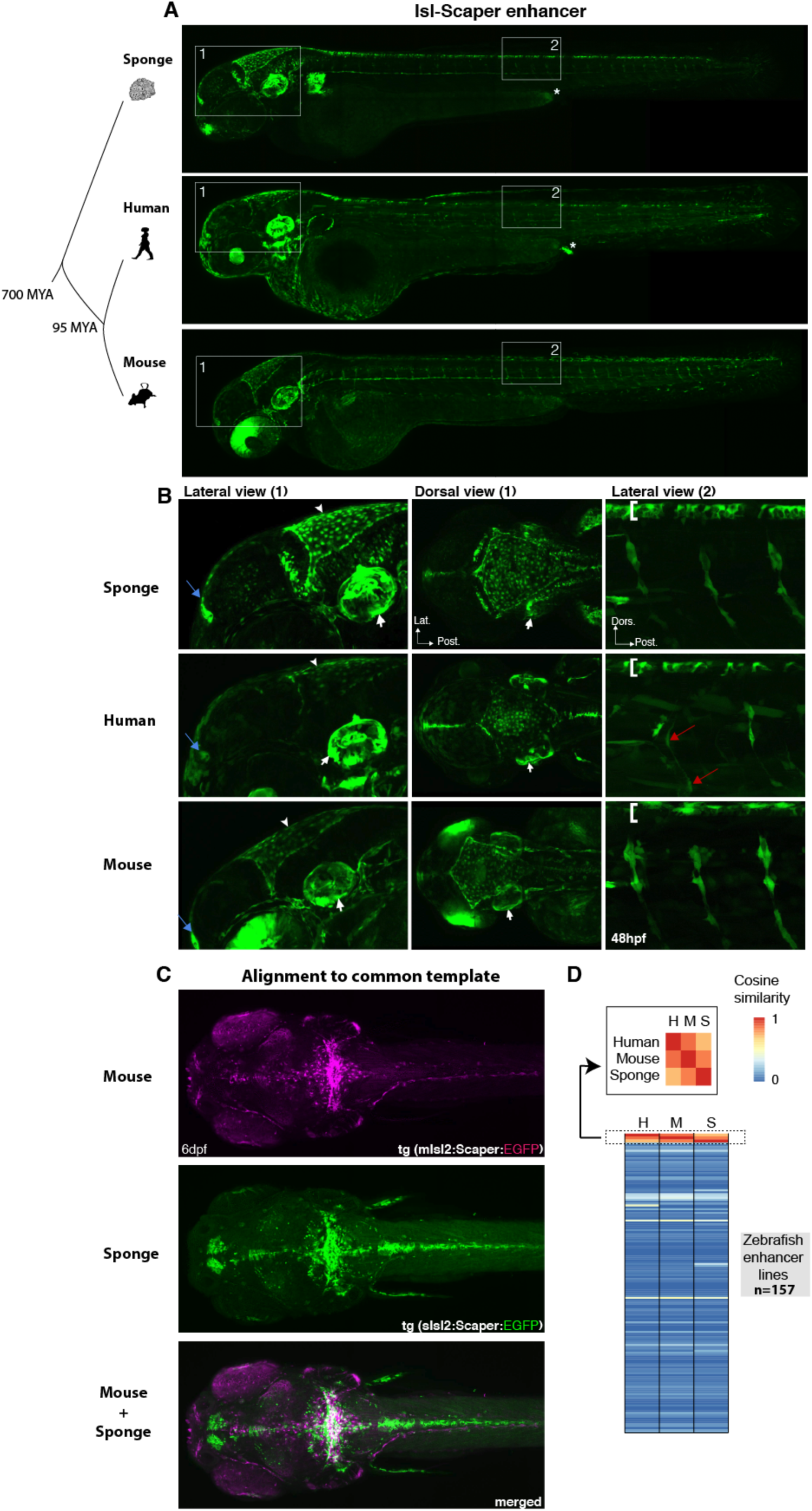
Mammalian homologs of sponge enhancer drive similar developmental expression. **(A)** Comparison of GFP expression pattern at 48 hpf across 3 stable transgenic zebrafish lines harbouring the sponge islet enhancer tg(*sIsl2-Scaper:egfp*) or corresponding islet ortholog enhancer tg(*hIsl2-Scaper:egfp*) tg(*mIsl2-Scaper:egfp*). Similar activity profile was detected in neural tissue, otic vesicle, pineal region and pectoral fin for the 3 ortholog enhancers. Both sponge and human enhancer shows activity in the proctodeum (asterisks). **(B)**. Close up of lateral and dorsal views from inset 1 and 2 in (a). Higher magnification reveals similar GFP expression in the neuro-epithelial layer of the hindbrain (lateral view (1), white arrowheads), otic vesicle (lateral view (1) and dorsal view (1), white arrows), pineal region (lateral view (1), blue arrows) pectoral fins (dorsal view (1), pink arrows) and in neurons of the roof plate (lateral view 2, bracket). Specific activity of the human enhancer is observed in the lens (lateral view (1), red arrowheads) and in endothelial cells (lateral view (2), red arrows), whereas the mouse enhancer drives GFP expression in the retina (lateral view (1), yellow arrowheads). (**C**) Neuroanatomical visualization of the mouse and sponge Islet:Scaper enhancers activity at 6 dpf using a common template built from confocal images. Subsets of neurons in the caudal part of the cerebellum are active in mouse and sponge enhancers (white signal in ‘merged’). **(D)** Heatmap comparison of anatomical locations showing reporter expression between sponge (‘S’), human (‘H’) and mouse (‘M’) *eISL* enhancers and 154 zebrafish enhancers shows greater similarity between *eISL* homologs (p<6.5e-3) (157 total lines compared). Inset shows *eISL* homologs only. See **FIG S17B** for full similarity heatmap.

To quantitatively assess the similarity of reporter expression between species, we compared against tissue-specific expression of 154 zebrafish enhancers from an enhancer trap assay at 88 anatomical locations (*40*). Similar to our transgenic lines, each enhancer is active in at least one type of neuron. For each line, we scored the presence or absence of GFP expression at the locations described by the enhancer trap experiment, with the inclusion of the fin as an additional category. We inversely weighted information theory scores to reduce the contribution of common description of expression terms, under the assumption that matched expression patterns from less common tissue types provided stronger support of a shared regulatory network (**Methods**). We calculated the cosine similarity for all pairs of enhancers using the weighted scores. Sponge, human and mouse enhancers shared activity in the same tissue significantly more often than expected (empirical p<6.5e-3)(**FIG4D, S16, S17)**. Among *Islet* enhancers, as expected, the mouse and human enhancers shared the most similar expression profile (mouse:human=0.86). Sponge and mouse enhancers shared more tissue-specific activity than sponge compared to human (sponge:mouse=0.83, sponge:human=0.71).

## Discussion

Although sponges and vertebrates diverged around 700 million year ago, regulatory enhancers from sponges drive cell type specific expression in zebrafish and mice. The sponge sequences lack discernable sequence similarity to vertebrate genomes, and sponge enhancer activity in vertebrates is not restricted to conserved chromosomal regions (i.e. microsyntenic units) and includes intragenic enhancers within sponge-specific genes. These results suggest that metazoans not only share common set TF families but also a suite of enhancers with a common regulatory grammar that can be interpreted across different phyla despite immense variation in body plans and developmental systems.

Sponge enhancers can direct similar cell-type specific expression patterns compared to mammalian sequences, thus these appear to be deeply conserved homologous enhancers. For enhancers to be independently maintained in disparate animal lineages, non-neutral selective forces must be operating on the elements across deep evolutionary time. The presence of common TF binding motifs in homologous enhancers suggests that these motifs may be critical for the maintenance of enhancer functionality. However given the absence of strongly conserved cis-regulatory motif order, selection most likely acts to maintain a cooperative binding environment, rather than to strictly preserve the ordering of protein binding (*22, 41, 42*). For instance, the *Islet* enhancers in sponges and vertebrates share a set of TF binding motifs that include Foxd1, Sp-1, Ap-1, yet these differ in order and frequency between homologs. Nonetheless, a core set of TF-DNA interactions may stabilize the enhancer over evolution, providing a foundation for further lineage-specific elaboration via the integration and dissociation of other *cis*-elements.

Regulatory co-option is an essential element of organismal evolution. By assessing sponge enhancers in the context of vertebrate development we reveal the nature of enhancer functional evolution in multicellular animals. Sponge development and body plans have little in common with vertebrates. Sponges lack muscles, nerves and a gut. Their cell lineages do not appear to progressively restrict to a germ layer and they lack a mesoderm; it is questionable as to whether they even gastrulate (*43*). Thus the expression patterns generated by the sponge enhancers in fish and mammals, which is largely restricted to neurectoderm lineages, suggests that gene regulatory circuits can be co-opted to perform lineage-specific developmental and cell type gene network functions (*44*). Given the apparent expansion of the number of enhancers in bilaterians after diverging from sponges (*1, 45*–*47*), which appears to parallel increases in metazoan TF family members (*48*), the duplication and divergence of ancestral enhancers would have yielded a larger repertoire of enhancers in vertebrates that, in turn, could be continually co-opted into new functions. We posit that the differential expansion of TFs and enhancers, and their subsequent co-option into new regulatory roles in eumetazoans underlies the evolution of this clade’s disparate complex body plans compared to the simple and restricted body plan of sponges.

## Acknowledgements

The authors would like to thank Gemma Richards, Ivy Chang and Emmanuelle Frampton for technical assistance and David Garfield for provision of data.

## Funding

This research was supported by Australian Research Council grants (DP160100573 and FL110100044) to BMD, National Health and Medical Research Council (NHMRC) grant (APP1107643) and NHMRC Career Developmental Fellowship (AP1111169) to MF, and Australian Research Council Early Career Award (DE160100755), a University of Queensland Early Career Grant (UQECR1832697) to EW. The Australian Regenerative Medicine Institute is supported by grants from the State Government of Victoria and the Australian Government.

## Author contributions

EW, FG, MF and BD were involved in the design of the experiments. ST and VG performed wet lab experiments. EW performed dry lab experiments. EW, ST, MF, BD, ES, GV and EM were involved in data analyses. EW wrote the manuscript, which was revised by MF and BD. EW, ST and MF were involved in figure construction. All authors commented on the manuscript.

## Data and materials availability

Sequencing data generated in this study can be found on ArrayExpress under E-MTAB-7846 (username: Reviewer_E-MTAB-7846; password: GA470bin). Other data generated from this work is available in the Supplementary Data File.

## Competing interests statement

The authors do not have any competing interests.

