## Supplementary material for "Early origin and deep conservation of enhancers in animals": sup_material

### METHODS

#### *Genome resources*

The following versions of genome assemblies were used: *A. queenslandica* (Aqu1), mouse (GRCm38/mm10), human (GRCh38/hg38), zebrafish (zv10) and fly (BDGP6/dm6).

#### *Amphimedon gene assignments*

Sponge transcripts were assigned human orthologs using a reciprocal best-hit BLAST [1] approach with human protein sequences. Treefam annotations [2] were also used to supplement orthology assignments. TFs were manually curated using similarity searching of genomic and transcriptomic sequence databases and literature searches. Additional methods were used for annotating TFs. These included using a Markov clustering algorithm to group similar transcripts (orthoMCL) and searches against TF protein domains from the DBD database of transcription factor predictions using hidden Markov models [3–5]. We further curated a set of TFs with robust orthology assignments for additional statistical tests to ensure our results (using all probable TFs) were also supported by TFs annotated with high confidence. These comprised of genes assigned by at least two of the above annotation methods (n=172).

#### *Identification of sponge enhancers based on histone marks*

We used chromatin immunoprecipitation sequencing (ChIP-seq) data sets for histone modifications in larval and adult *Amphimedon* where the most common combinations of regulatory states were determined by a classification algorithm [6]. We intersected the enhancer regions (avoiding the 'weak enhancer' state) to genome positions of previously defined conserved microsyntenic pairs (overlap 1 bp) [7]. Genomic locations of sponge enhancers present in conserved syntenic regions are provided in **SFile**.

#### Identification of homologous enhancers by TF composition

To identify common motifs, we scanned both query and target sequences using position-specific scoring matrices (PSSMs) with LASAGNA [8]. The LASAGNA algorithm is based on the premise that motif from each class of transcription factor binding site (TFBS) is comprised of a short core sequence with flanking bases, which may vary in identity and length. We used PSSMs generated using this method with sequences from the TRANSFAC database to identify TFBS in our sequences and measure motif significance using a probabilistic zero-order Markov model constructed from the base frequencies for each motif.

Next, we took sliding windows of the query sequence, where each 'windowed' sequence is of a similar length to the target enhancer (~1kb). All motifs below a default PSSM scores cut-off of $p$=0.01 were retained and the frequency of each motif was tabulated for each sequence. To improve specificity, we further restricted motif significance to a more stringent threshold for the sponge query sequence (p=1e-3) under the assumption that the sponge sequence may provide the molecular handle to retrieve the divergent sequences.

Each sequence, target and windowed query, was normalized to unit length (L2 normalization) to make comparable sequences of different lengths, and weighted by target motif significance by multiplying against the maximum PSSM score for each motif in the target sequence.

Only motifs common between target and query sequences are then compared and a similarity score calculated by cosine similarity:

$$\cos\theta = \frac{\vec{a} \cdot \vec{b}}{\|\vec{a}\|\|\vec{b}\|}$$

The R text-mining package 'tm' was used to facilitate data import, sequence handling, and the creation of motif score matrices.

***Comparison of expression similarity with enhancer trap lines***

To quantitatively assess the similarity in GFP expression patterns between species, we compared the anatomic descriptions of enhancer activity from 154 stable zebrafish lines from a Gal4 enhancer trap (ET) screen with matched descriptors from our fish lines [9]. The ET transgenesis method involved the random insertion into the genome of a transgene linked to a minimal promoter, which when inserted near an endogenous enhancer element expresses GFP in a pattern determined by the enhancer. The majority of lines, screened by reporter expression in founder, have expression in the central nervous system (CNS)(82%)[10]. The experiment preferentially raised F1 larvae where at least one tissue showed neuronal expression at 3-5 dpf. Fittingly, all stable lines generated in this study showed neuronal expression patterns.

Our fish were described in a similar manner (confocal microscopy) at a comparable time point to the published dataset. Descriptions were filtered to reduce subjectivity in descriptions and to generalize complex expression patterns. Specifically, we classified expression in the eye into either partial or total. More subjective descriptors such as ‘weak’, ‘strong’, ‘sparse’ and ‘bright’ were removed. We then used the inverse of Shannon’s entropy measure to reduce the contribution of common description of expression terms. Indeed, certain patterns (particularly in pineal, muscle and heart) suggested as possible background expression independent of the endogenous enhancers are thus accounted for using the weighting scheme.

Using the catalog of where GFP was active in each enhancer trap fish line, we constructed a binary locational matrix for each enhancer line and inversely weighted each anatomical location term by their frequency across all lines, such that the most frequent regions was given the lowest weight (**Fig S18)**. We characterized the *Islet* stable lines in this study (sponge, mouse, human) at a comparable developmental time (72 hpf), by similar ventral and dorsal imaging with a confocal microscope. Pairwise similarity measures were calculated by cosine similarity. A summary of all anatomical description expression patterns and full similarity matrix is provided (**SFile)**.

***Statistical test for conservation of motif order***

To increase the sensitivity in detecting conservation of motif order, we accounted for evolutionary divergence by filtering PWMs to merge common gene families. This was achieved by removing numbers following TF gene genes. After accounting for lineage-specific gains and losses, we identified 37 common motifs among the orthologous sequences from four species (sponge, zebrafish, human and mouse). We assigned a unique character to each motif and used the Needleman-Wunsch [11] and Smith-Waterman [12] algorithms to solve for global, global-local and local alignments (implemented in the R package ‘Biostrings’ v2.40.2). We tested for conservation of motif order between all sequences in a pairwise manner in both forward and reversed directions. We generated empirical null distributions to assess significant similarity by scrambling the motif order for the target sequence while keeping the query sequence and other parameters the same, and recording the top alignment score 100 times.

***Transgenic zebrafish reporter assays***

The transgenic lines, including *Tg(sisl2-scaper:eGFP)*$^{uq5mf}$, *Tg(sccne1-c19orf2:eGFP)*$^{uq6mf}$, *Tg(stdrd3-diaph3:eGFP)*$^{uq7mf}$, *Tg(hisl2-scaper:eGFP)*$^{uq8mf}$ and *Tg(misl2-scaper:eGFP)*$^{uq9mf}$ were generated using Gateway cloning (Life Technologies). Briefly, these constructs were generated by PCR from sponge, human and mouse genomic DNA, respectively using the primers described in **SM**, cloned via Gateway pME vector (pDONR221) into the Zebrafish Enhancer Detector (ZED) vector [13]. To generate the transgenic strains, 70 ng/µL of plasmid DNA and 25 ng/µL of *tol2* transposase mRNA were injected in one-cell-stage wild type zebrafish embryos. An enhancer sequence of interest was cloned upstream of *gata2* promoter and *gfp* transgene with an internal control (RFP) for transgenesis efficiency and GFP normalization. The internal control is driven by cardiac actin promoter and is expressed in muscle cells. GFP and RFP positive embryos ('double positive') were selected at 24 hpf based on GFP brightness and grown for 3 months until sexual maturity. In total, around 30% of $F_0$ fish showed germline transmission (**Table S1)**. These fish were then crossed to wild type to generate transgenic stable lines (**Table S2**). Similar reporter expression patterns were observed between GFP positive F1 individuals from the same founding fish line. To resolve cases of multiple vector insertions, we outcrossed the line until the clutches show equal segregating ratios of GFP positive and GFP negative individuals.

Zebrafish were maintained at the University of Queensland under standard husbandry conditions with a 14-hour light and 10-hour dark cycle. All animal experiments were performed in accordance with the guidelines of the animal ethics committee at the University of Queensland (IMB/237/16/BREED). Primers used are in **Table S3**.

***Generation of mouse transient lines***

All procedures were performed in accordance with the Australian Code of Practice for the Care and Use of Animals for Scientific Purposes. Experiments were approved by the Monash Animal Ethics Committee, under project number 17657. Generation of mouse transient transgenics was performed by the Monash Genome Modification Platform (MGMP). Briefly, constructs were provided to MGMP for purification and linearization followed by pronuclear injections of each construct into fertilized mouse embryos. Embryos were transferred to recipient pseudo-pregnant females and embryos were collected at embryonic day (E) 10.5.

***Mouse embryo dissections and LacZ staining and sectioning***

Pregnant mice were euthanised and embryos were dissected away from the uterine horn and extraembryonic tissues in cold 1X PBS. The fourth ventricle was pierced to prevent any future trapping of staining. For LacZ staining, the Wellcome Trust Sanger Institute protocol was used (ftp://ftp.sanger.ac.uk/pub/resources/mouse/sigtr/XGalStaining.pdf). Embryos were immediately placed into 0.2% glutaraldehyde supplemented with 5 mM EGTA in 0.1M phosphate buffer (pH 7.3) for 30 min rocking at room temperature. Embryos were washed 3 x 15 min rocking at room temperature (RT) in 0.1M phosphate buffer (pH 7.3) supplemented with 2mM $MgCl_2$, 0.01% deoxycholate and 0.02% NP-40 (wash buffer). Following washes, embryos were transferred to stain solution (wash buffer supplemented with 5 mM potassium ferrocyanide, 5 mM potassium ferricyanide and 1 mg/ml X-gal) rocking at 37°C overnight. If staining came up quickly, within 1-2 hours then staining for those embryos was stopped. Following LacZ staining, embryos were washed 3 x 15 min in wash buffer while rocking and fixed in 4% paraformaldehyde for 30min at RT. After the fixation, embryos were washed with 1X PBS 3 x 5min rocking at RT. Embryos were stored at 4°C or embedded. Whole embryos were imaged on a black background with a Zeiss SteREO Discovery V20 microscope.

For gelatin embedding, embryos were treated 5% and 20% sucrose in 1X PBS until the embryos sank. Embryos were incubated in 7.5% gelatin/15% sucrose for 2 hours at 37°C. Embryos were cut into three pieces (to obtain the best cross-sections), embedded in gelatin on ice and transferred to dry ice for freezing once the gelatin was set. Embryos were sectioned on the Leica CM 3050S cryostat at 20 µm. Gelatin was removed from sections by incubation with 1X PBS at 37°C for 10 minutes. Sections were fixed with 4% PFA for 10 min and washed 3 x 5min with 1X PBS. Slides were mounted with fluoromount-G (Southern Biotech) and imaged on the Zeiss Axioplan Imager Z1.

***Imaging and characterization of transgenic fish expression***

Live and fixed embryos were mounted laterally or ventrally in 1% low-melting agarose and imaged using a Zeiss LSM 710 FCS confocal microscope (Zeiss, Germany). Images were taken under 20x objective lens (using z stacks 4-5 µm apart) or 40x objective lens (using z stacks 1 µm apart). Images in this paper are maximum projections of z series stacks. Images were processed using ImageJ (National Institute of Health) [14]. Anatomical locations of GFP expression was recorded at three time points 24, 48 and 72 hpf.

***Confocal overlay to common template***

We used the Advanced Normalization Tools (ANTs, github.com/ANTsX/ANTs) to build a common template from our confocal images [15,16]. The DAPI channel stacks were used to build the template, using the antsMultivariateTemplateConstruction2 command with 30 iterations and a gradient step of 0.02. The resulting warps were applied to the remaining channels of the confocal stacks to align all of the expression patterns in the same frame of reference.

#### *Fluorescent in situ hybridisation and immunofluorescence staining*

Whole mount fluorescent *in situ* hybridisation (FISH) in zebrafish embryos was performed as previously described [17,18]. Probes used were *gfp* [19] and *Isl2a*. The *Isl2a* probe was generated by PCR from zebrafish embryo cDNA pool at 5 days post fertilisation (dpf) and reverse transcribed with T7 polymerase (New England Biolabs). The *Isl2a* probe was labelled with Tyramide Signal Amplification (TSA) Plus Cy3 Solution (1:100, Perkin Elmer). Whole mount immunofluorescence staining for GFP was performed as described [20]. The following antibodies were used: chicken α-GFP (1:250, Abcam) and goat anti-chicken IgG Alexa 488 (1:400, Invitrogen).

Primers for wholemount zebrafish FISH:

Isl2a_ISH_F: CCGATTAGACATGACACCACAG

Isl2a_ISH_R: TGTAATACGACTCACTATAGGTATGGACGGTTGTCCTGA (T7 promoter = underlined)

#### *Fluorescent Activated Cell sorting (FACS)*

72hpf zebrafish embryos were anesthetised with tricane diluted 1:1 in the embryo medium and the yolks were mechanically removed by pipetting up and down at least 10 times in calcium free ringer's solution. Zebrafish embryos were further digested in PBS containing 0.25% liberase at 28°C for 5 min and a single cell solution was prepared by passing through a 40μm nylon mesh. Both GFP positive and negative single cells were sorted into Trizol LS reagent (Life Technologies), respectively with a BD FACSAria Cell Sorter (BD Biosciences) (**Table S4**). Subsequently, RNA was extracted using a Direct-zol RNA extraction kit (Zymo research).

#### *Smart-Seq2 library preparation and sequencing*

RNA-Seq libraries were prepared from purified total RNA using a modified Smart-Seq2 protocol developed by Picelli *et al.* [21]. 2 ng of purified total RNA (0.4 ng/μL) was combined with 1 μL of 10 μM oligo-dT primer (/5Biosg/AAGCAGTGGTATCAACGCAGAGTACT$_{30}$VN; Integrated DNA Technologies) and 1 μL of dNTP mix (10 mM each; Invitrogen, y02256), then the protocol was continued as described (ref. 2). Briefly, the RNA was reverse transcribed with the Smart-Seq2 TSO (/5Biosg/AAGCAGTGGTATCAACGCAGAGTACATrGrGrG, Integrated DNA Technologies), followed by 12 cycles of PCR amplification to obtain enough cDNA to prepare a library. Volumes of reagents were scaled accordingly to maintain final concentration ratios as in the original protocol, except for the PCR preamplification where the Smart-Seq2 ISPCR primer (/5Biosg/AAGCAGTGGTATCAACGCAGAGT; Integrated DNA Technologies) was added to a final concentration of 0.25 µM. 0.5 ng of cDNA was prepped into a library using the Nextera XT DNA Library Prep Kit (Illumina, FC-131-1096), with 12 cycles of PCR used to amplify the final library. The final Nextera XT libraries were quantified on the Perkin Elmer LabChip GX with the DNA High Sensitivity Reagent kit (Perkin Elmer, CLS760672). Libraries were pooled in equimolar ratios.

Sequencing was performed using the Illumina NextSeq500 (NextSeq control software v2.1.0 / Real Time Analysis v2.4.11). The library pool was diluted and denatured according to the standard NextSeq protocol (Document # 15048776 v05), and sequenced to generate single-end 76 bp reads using a 75 cycle NextSeq500/550 High Output reagent Kit v2 (Illumina, FC-404-2005). After sequencing, fastq files were generated using bcl2fastq2 (v2.18.0). Library preparation and sequencing was performed at the Institute for Molecular Bioscience Sequencing Facility (University of Queensland).

***Transcriptomic analyses of GFP sorted cells in stable fish lines***

Transcriptomes for FACS-sorted GFP positive and negative cell population produced with the Smart-Seq2 protocol was assessed for differential gene expression. All comparisons were performed in duplicates and each sample required pooling of cells from multiple fish of the same genetic background to improve the numbers of GFP positive cells (in particular *eCCNs*)(**TableS4**).

Reads were mapped to the zebrafish genome assembly (GRCz11) using Bowtie2 software [22]. The RSEM program [23] was used to quantify gene and transcript abundance. A second method of read mapping, was used to measure transcript abundance by mapping to the zebrafish transcriptome (Ensembl Release 95) using the Salmon software on quasi-mapping mode [24]. Both methods produced comparable count matrices. As expected, GFP was consistently highly enriched in the GFP positive samples (>2,000 counts) and fewer than 10 reads, on average, were detected in GFP negative samples. For downstream analyses we used the RSEM/Bowtie2 counts matrix. Differential expression analysis was performed using the R packages 'limma' and 'edgeR' [25,26]. Significance was called using empirical Bayes moderated-t p-values (FDR<0.1) relative to a minimum log2 fold-change of 1 to select for changes of large effect size, as these were more likely to be biologically significant. DAVID [27] was used for functional enrichment analyses. For both differential expression and functional term enrichment analyses, p-values were adjusted for multiple testing with the Benjamini and Hochberg method [28].

### SUPPLEMENTAL TABLES AND FIGURES

***Table S1. Numbers of injected embryos and $F_0$ founders***

| Enhancers | Injected embryos | Double positive embryos | Percent transformed |
|---|---|---|---|
| *eISLs* | 33 | 8 | 0.24 |
| *eCCEs* | 47 | 10 | 0.21 |
| *eTDRs* | 27 | 8 | 0.30 |
| *e1s* | 18 | 9 | 0.50 |
| *e2s* | 30 | 9 | 0.30 |
| *eIShs (hs)* | 31 | 12 | 0.39 |
| *eISLm (mm)* | 66 | 10 | 0.15 |

***Table S2. Numbers of GFP positive transgenic zebrafish following outcrossing***

| Enhancers | Number of zebrafish | | |
|---|---|---|---|
| | F1 generation | F2 generation | F3 generation |
| *eISL* | 15 | 29 | 21 |
| *eCCN* | 16 | 38 | - |
| *eTDR* | 38 | - | - |
| *e1s* | 6 | - | - |
| *e2s* | 3 | - | - |
| *eIShs (hs)* | 3 | 46 | - |
| *eISLm (mm)* | 11 | 2 | 70 |

***Table S3. Primer sequences for cloning into vectors for transgenesis***

| | Sequence (5'-3') |
|---|---|
| *sisl2-scaper*_F | TTTTGGCTTGATTGCTGTTG |
| *sisl2-scaper*_R | ACCTGCCCATACACACAGTC |
| *sccne1-ccne1*_F | CCCTTGTGTCCTCCACTTTC |
| *sccne1- ccne1*_R | TCTGCTGTGCCTCTCTAACC |

| | |
|---|---|
| *stdrd3-diaph3*_F | GGGAATAGCAAATCCAGGAC |
| *stdrd3-diaph3*_R | TTGGATTTGGAACAAACAGC |
| *e1s* Aqu2.1.24542_001_F | GATTCTGTGGACCCATGCTT |
| *e1s* Aqu2.1.24542_001_R | GAGAGACAGCCATGCAACAA |
| *e2s* Aqu2.1.25069_001_F | CGCACCTTCTTGACCTTCAG |
| *e2s* Aqu2.1. 25069_001_R | CCGAGATAGAGGAAGAGAAACTC |
| *Human_sl2-scaper*_F | **GGGGACAAGTTTGTACAAAAAAGCAGGCT**TCTCCTTGGTGAGGGCATTTAG |
| *Human_isl2-scaper*_R | **GGGGACCACTTTGTACAAGAAAGCTGGGT**CACACATGCACAGTAAGTTAAGCC |
| *Mouse_isl2-scaper*_F | **GGGGACAAGTTTGTACAAAAAAGCAGGCT**TCTTGGGGATCTTCTTTCAGGA |
| *Mouse_sl2-scaper*_R | **GGGGACCACTTTGTACAAGAAAGCTGGGT**CTTCCTCCCTCATAAGACACAAGT |
| *zfisl2-scaper_F* | **GGGGACAAGTTTGTACAAAAAAGCAGGCT**GTACAGCATAGCGTACTTTGTGATT |
| *zfisl2-scaper_R* | **GGGGACCACTTTGTACAAGAAAGCTGGGT**ATTCAGTGGGACCCATTTGTGG |

- Gateway homology arm = underlined and bold

***Table S4. Number of GFP positive cells by FACS sorting***

| Enhancers | GFP positive cell numbers | No. of fish embryo | No. of events/fish embryo |
|---|---|---|---|
| *eISLs* | 10000 | 143 | 70 |
| *eCNNs* | 1082 | 72 | 15 |
| *eTDRs* | 50000 | 100 | 500 |

### SUPPLEMENTAL RESULTS

#### *Transcriptomic comparison of GFP sorted cells in stable fish lines*

Similar numbers of genes were found expressed across all three lines. On average, approximately 19,500 genes were used for expression analyses after filtering out those with low counts (below 10). The number of differentially expressed genes between GFP positive and GFP negative fractions varied between enhancers. Comparative analyses of the transcriptomes revealed a substantial number of genes differentially regulated in the *eISLs* and *eTDRs* fish lines (360 and 1702, respectively; FDR<0.1) (**Methods**)(**FIGS6**). In the *eCNEs* line, 10 differentially regulated genes were identified (FDR<0.1). The low number potentially reflects the low number of gpf positive cells harvested (**TABLES4**).

Additional to the functional term enrichment analysis described for eISLs – for the eTDRs transgenic line, upregulated genes were associated with neuron differentiation, nervous system development and axonogenesis (p.adj<2e-3), consistent with observed pan-neural reporter expression. In addition, genes linked to multicellular development were also upregulated. These genes involved the establishment of localization and regionalization, somitogenesis, segmentation and anterior/posterior pattern specification (p.adj <0.05). In the eCNEs fish line, a slight enrichment for neural crest cells was detected supporting the location of enhancer active cells in the head and spinal cord (p<0.1).

#### *TF motif composition of sponge enhancers*

We characterized TF binding motifs by aligning 398 Transfac PWMs to sponge enhancer sequences using the LASAGNA (Length-Aware Site Alignment Guided by Nucleotide Association) algorithm [8]. At a p-value<1e-3, 117, 160 and 131 motifs were identified for *eISLs*, *eTDRs* and *eCCNs*, respectively. We compared motif composition across the sponge enhancers and found 44 motifs common across the sequences, with 22, 34 and 20 motifs unique to *eISLs*, *eTDRs* and *eCCNs*, respectively (**SFile**).

#### *TF binding motifs discriminate TFs that drive expression of Islet in* **Amphimedon**

To show that sponge enhancer likely drive Islet expression and that TF binding motifs are predictive of the collaborative TF binding landscape, we test the relationships between genes based on transcriptomic data in the sponge. We theorized the following 1) TFs recruited to an enhancer should show co-regulated gene expression with genes that are directly regulated by the enhancer; 2) the collective of TFs recruited to an enhancer that drives expression can be estimated by the TF binding motifs present on the enhancer. We can separate sponge TFs into two groups: 1) genes presumed to bind the regulatory element based on the presence of binding motifs on the enhancer ('enhancer TFs') 2) genes that cannot be associated by motif to the enhancer ('non-enhancer TFs'). In this way, a significant relationship between sponge Islet and enhancer TFs not only suggest that Islet is regulated by the enhancer but also that the enhancer is driven by a collection of TFs, which can be predicted by binding motifs.

We first characterized the TF motif composition of the sponge enhancer using Transfac position weighted matrices (PWMs) and identified a total of 119 motifs on the putative sponge Islet enhancer sequence (p-value<1e-3). We filtered out motifs denoting protein-protein interactions and matched annotated sponge TFs to candidate TF binding motifs. 30 (of 525) sponge TFs were associated to motifs on the sponge enhancer. Where two sponge TFs could be assigned to a single binding motif, only one sponge TF was randomly chosen. To be as precise as possible, we did not associate TF binding motifs to cases where sponge TFs can only be annotated at the broad level of TF class.

To infer regulatory relationships between genes, we used a dense CEL-seq time-course across embryonic development [29]. Using Pearson's correlation coefficient, we tested the relationship of Islet gene expression to TFs associated to the enhancer versus Islet with non-associated TFs (**FIG 1f**). We found a significant increase in Pearson's correlations between Islet and those TFs with motifs on the enhancer compared to TFs without linked motifs on *eISLs* (Mann-Whitney U, p=5.5e-4). Reducing the genes studied to a high confidence set of 172 TFs did not strongly influence the magnitude of fold change observed. We observed a log2 fold change of -4.8 for all annotated TFs compared to -3.5 when restricting the analyses to TFs annotated with high confidence.

We further found a difference in the relationship of the TFs with the Islet adjacent 'bystander' gene Scaper. Scaper expression was less correlated with the *eISLs* motif associated TFs than Islet (Mann-Whitney U, p<0.05) suggesting the regulatory element is more likely to regulate Islet than Scaper. FDR was used to adjust p-values in multiple testing. Together, these results show the annotation of sponge sequences with PWMs is able to detect the subset of TFs that drive expression of Islet in *Amphimedon*.

To assess our background, we tested for correlations between the microsyntenic enhancer-associated TFs and randomly selected genes (n=100) and found their correlation coefficients to be low (median

Pearson's rho=0.02). To roughly assess the impact of expression level, we further restricted the comparison to highly expressed genes and found similarly weak linear relationships (defined as genes of mean expression > 50)(median Pearson's rho=-0.01). Finally, unlike the genes tested in our study, there was no discernable difference between correlations using TFs linked to enhancers and un-associated TFs when compared to randomly selected genes.

In sum, although it was not possible to infer a sponge ortholog for many PWMs and many TFs do not bind recognizable motifs, PWMs appear able to identify TFs on sponge enhancers that are correlated with genes in the microsyntenic unit.

***Sponge-eumetazoa alignments***

We performed whole genome alignments of the sponge genome against human, mouse, zebrafish and fly genomes from Ensembl. We used blastn to align repeat-masked eumetazoan genomes and a relaxed expectation value (E-value) cutoff of 1e-3 to allow for sequence divergent hits. All hits that met this threshold were kept and overlapped with functional annotation regions. Results are summarized in **FIGS1c**.

Although genome-wide alignments are unable to identify a candidate human ortholog to sponge Islet (source: UCSC genome browser), we investigated several alternative alignment algorithms including more sensitive approaches. These strategies included first reducing the sequence search space by restricting the region to only encompass the span of Scaper and using blastn [1], a heuristic alignment approach which is more sensitive than the genome-wide alignment algorithms (database=human refseq genomic, default parameters). To optimize sensitivity, we also experimented with reducing the word size (word size={4, 11}). We further used an algorithm guaranteed to identify the optimal match, the Smith-Waterman dynamic programming algorithm [12] implemented in EMBOSS Water [30], to perform a local alignment between the sponge and human sequences using different parameters (gap open={5, 10} and gap extend={0.1, 0.5}). Neither the heuristic searches nor the optimally aligned region by Smith-Waterman could recover the human region identified using the TF motif composition method, which mapped to the mouse enhancer identified using mouse ENCODE data [31] and exon synteny information (see below).

***Identification of mammalian orthologs to sponge enhancers based on TF composition at conserved syntenic regions***

We used the method to interrogate conserved syntenic regions and human and mouse using candidate sponge enhancer sequences. We assessed the accuracy of the method in two ways. First, we looked for overlap with available functional genomics information. For example, we used mouse ENCODE data to infer enhancer activity based on histone marks in specific tissues. Second, we assumed the identified regions are good candidates for the sponge enhancer if the human and mouse regions were also reciprocal best-aligned regions by whole genome alignment (source: UCSC Genome Browser). Mouse ENCODE datasets used were of narrow peak format and only peaks consistent between two biological replicates were used.

*Isl2-Scaper*

The candidate sponge enhancer (709bp) was used to identify candidate homologs in mammals. We first performed the search against the human genomic sequence spanning the Scaper gene (chr15:76347904-76905444), a region of ~558kb. A window size of 900bp was used. The top hit was a region located at chr15:76358695-76359594 (score=0.87). This region is located in the 3' end of Scaper, similar in location to the enhancer in zebrafish [7]. Although the region cannot be aligned to the zebrafish genome, the sequence was mappable to the mouse genome (UCSC Genome Browser). Based on the tissue-specificity of the mouse histone mark H3K4me1 during embryonic development (ENCFF053GHW (face), ENCFF191NER (midbrain), ENCFF350UJK (forebrain)) and the relative location of the histone modification based on conserved exon structure of Scaper between mouse and zebrafish, this mouse region appeared a likely candidate mouse ortholog of the sponge sequence. We also applied the algorithm to search for the Isl2 enhancer in mouse using the sponge enhancer as query. The region (chr9:55549879-55938119) spanning the entire Scaper gene (~388kb) was compared to the sponge enhancer with a bin parameter of 1kb. In this case the 5$^{th}$ ranked hit (chr9:55561854-55562852) overlapped the region predicted by the histone marks.

No conservation with respect to gene structure and Scaper transcripts was found between sponge and vertebrate Isl/Isl2 enhancers. Although Scaper protein sequences can be aligned between sponge and mouse Scaper, the location of these enhancers with respect to the Scaper gene structure was different and did not overlap. Indeed, there was no discernable similarity in exon spacing between sponge and eumetazoan genes. To further investigate, we examined the genomic location of the enhancer relative to the protein sequence of Scaper and compared this between sponge and mouse. We did not identify a correspondence between sponge and mouse. In contrast, both predicted mouse and fly enhancers were located in exons close to the 3' end of their longest Scaper transcript, although the protein sequences in this region were too divergent to be robustly aligned (blastp).

*Tdrd3-Diaph3*

The mouse genomic sequence for Diaph3 spanning 490kb was extracted (chr14:86657341-87147735). Using the candidate sponge enhancer for Tdrd3 located in Diaph3 (Contig13428:40966-42475) marked by H3K4me1 and H3K27ac, we used a bin parameter of 1.5kb to identify similar regions in the mouse sequence based on TF binding motif composition. A total of 247 regions within the mouse Diaph3 gene were searched and the best scoring hit was found at chr14:87141745-87143240 (score=0.89). The region partially overlapped with H3K4me1 marks in the midbrain, forebrain and face in mouse at E10.5, consistent with pan-neuronal GFP activity in sponge Tdrd3 enhancer transgenic but also H3K4me3, suggesting this may be a promoter rather than an enhancer. Based on its location at the 5'end of mouse Diaph3, we extracted 169kb of human sequences including approximately 5kb of sequence immediately upstream of Diaph3 from the human genome. Using the same parameters as used in mouse, we interrogated 114 regions. While best hit at chr13:60110151-60111639 (score=0.81) did not match the best mouse hit, the human orthologous sequence to the best scoring mouse region was located in a highly scored region (chr13:60165226-60166715; score=0.79; rank 5). We adjusted the method to account for background motif composition by down-weighting common motifs and used a randomization procedure to calculate a p-value (described below). This had the effect of identifying nine regions with significant similarity to the query sponge sequence. Notably, the top ranking hit after adjustment for genome-specific background motif composition was the human region (chr13:60165226-60166715) that best aligned to the candidate mouse homolog. Although accounting for background motif signals produced consistent results for the Tdrd3 enhancer, this method did not output reciprocal best-hit results for mouse and human Isl2 enhancers. We further note without the down weighting of common motifs, 30 human regions with p.adj<3e-3 (score more significant than any permuted region) were identified. These regions can be mapped to 8 regional clusters of hits. In total, the hits span 21 loci. In mouse, 21 significant regions were identified. Such regions may represent homologous enhancers, but it will require functional tests to verify.

*Ccne1-Uri1*

Given the sponge enhancer tested in the microsyntenic region of Ccne1-Uri is located within the gene span of Ccne1 (931bp), we extracted the human and mouse region encompassing Ccne1 to the beginning of Uri (human: chr19:2803981-29917977; mouse: chr7:38018408-38109354) and tested their TF composition against sponge enhancer. A total number of 76 human and 95 mouse ~1kb genomic regions were tested. We took the top two highest scoring regions located at chr19:29854374-29855574 and chr19:29911968-29913168 with scores of 0.85 and 0.84, respectively. The first region appeared mammal-specific and aligned to a mouse region that overlapped H3K4me1 marks in midbrain, face and forebrain (E10.5), the second region is interspersed with SINE elements and did not align to genome sequences outside of primates. This region was located more much proximal to Uri1 than Cnne1. Similarly, the top hit in mouse was located in a repeat-rich region that appeared rodent-specific but overlapped H3K4me1 marks active in the forebrain and midbrain (E10.5) indicating enhancer function. Similar to the second human hit, this region was also located upstream to Uri. Given our sponge transgenic show reporter gene expression in zebrafish fore- and midbrain and syntenic genomic location with respect to the human hit, this mouse region could be an ortholog to the sponge enhancer.

Since the sponge enhancer tested is located within Ccne1, we further restricted the search region to the Cnne1 gene and its flanking region (5kb)(human: chr19:29807898-29828308; mouse: chr7:38095984-38109534). These regions were ~20kb and ~14kb, respectively. In mouse, we examined the top three hits. The first region (chr7:38108560-38109541) matched overlaps a TFBS for Cdx1 (Source: PAZAR; OREG1475323), while the second hit (chr7:38107592-38108560) overlaps a Spi1 TFBS (Source: Jaspar; OREG190146). Both these regions were located immediately upstream of Ccne1 and were enriched for H3K4me1 marks in brain and face at E10.5 (Source: Encode). The third ranked region chr7:38099854-38100821 was intronic and overlapped a Cdx1 binding site (Source: PAZAR; OREG1475322). In human, the first two top ranked regions were scored equivalently (0.84). One was located in an intron near the 3' end of the gene and encompassed a SINE element (chr19:29818584-29819556). The other region was also intronic (chr19:29813726-29814698), but located close to the 5' end of the gene. This region overlapped Encode ChIP-seq binding sites for GATA1, TAL1, GATA2, and a motif for SMARCA4 (Source: PAZAR; OREG1250838).

***Adjustments to TF composition alignment***

We made several adjustments to the described method that may improve the method's ability to discriminate between background and actual signal. These include an adjustment for background motif frequencies and a method to calculate a p-value based on the randomization of motif locations within a sequence. We note that the adjustment to include background TF motif frequency did not always produced consistent results (i.e. aligned regions in human and mouse to the sponge enhancers were not reciprocal best hits for human-mouse by whole genome alignments) and suggest that this is due to the underlying regulatory structure specific to each enhancer. The adjustments are described below:

**Background motif frequencies**

To adjusting for background motif frequencies, for each species, we randomly sample 500 sequences of length $l$, here 800bp was used, to define the baseline occurrence frequencies of TFBS in the respective genome. The average frequency of motif $i$ in genome $g$ is obtained:

$$m_{g,i} = \frac{\sum_{a=1}^{t} x_a}{t}$$

where $m$ is the arithmetic mean of motif $i$ in genome $g$. $x$ is a binary value denoting presence/absence of motif $i$ for a given sequence $a$ from genome $g$, $t$ is the number of random sequences scored.

Both query and target sequences were then weighted using the background motif probabilities. Any motif in either target or query, where the expected number of occurrence by chance is greater than one is down-weighted by multiplying by the inverse of the expectation of the motif (i.e. $1/m_{g,i}$) in the genome of interest.

**P-value calculation**

Finally, a randomization procedure is used to assign statistical significance by computing similarity scores where those motifs passing motif significance cut-off in the query sequence are randomized with respect to their genomic locations. We assigned a *p*-value by the following procedure – by giving a *p*-value of 0 if the similarity score is above the highest value of the null; a p-value of 1/n, where n is the total number of queried sequences, if between the first and second highest values of the null; and 2/n if between second and third and so on. The Benjamini-Hochberg method [28] was then used to correct for multiple testing.

***Statistical test for conservation of motif order***

We detected evidence of motif order conservation ($p<0.05$) by global-local alignment between sponge and human enhancers, and sponge and mouse elements ($p=0.02$ and $p=0.04$, respectively) but not between zebrafish and sponge (**SFile**). These alignments were short (2-3 motifs). On close inspection of the aligned regions, we found conservation at TF motifs from genes of the same TF class at overlapping locations, suggesting they may not reflect true conservation of binding order but rather PWMs from highly related genes from the same gene family. Furthermore, the best-aligned regions between sponge and human generally did not match the best-aligned regions in the sponge–mouse alignment (for alignments see **SFile)**. The zebrafish sequence aligned poorly with the mammalian sequences except when locally aligned with the human sequence ($p=0.01$).

We further asked whether less specific models based on TF classes/protein-binding domains could detect binding motif order. We matched the 37 shared motifs between the four sequences to 13 protein-binding domains (**FIG3g**). We found the best local match across all pairwise species comparisons to be variable. For example, the the optimal alignment between the sponge and zebrafish sequence comprised of two HSF domains, a C4 zinc finger domain, a bZIP domain and a C2H2 zinc finger domain (p=0.01), whereas between mouse and zebrafish a sequence containing an HSF, PAX, bZIP, PAX and zinc finger C2H2 was the best alignment (p=0.02).

The parameters used for local alignments were gap opening=5 and gap extension=2.

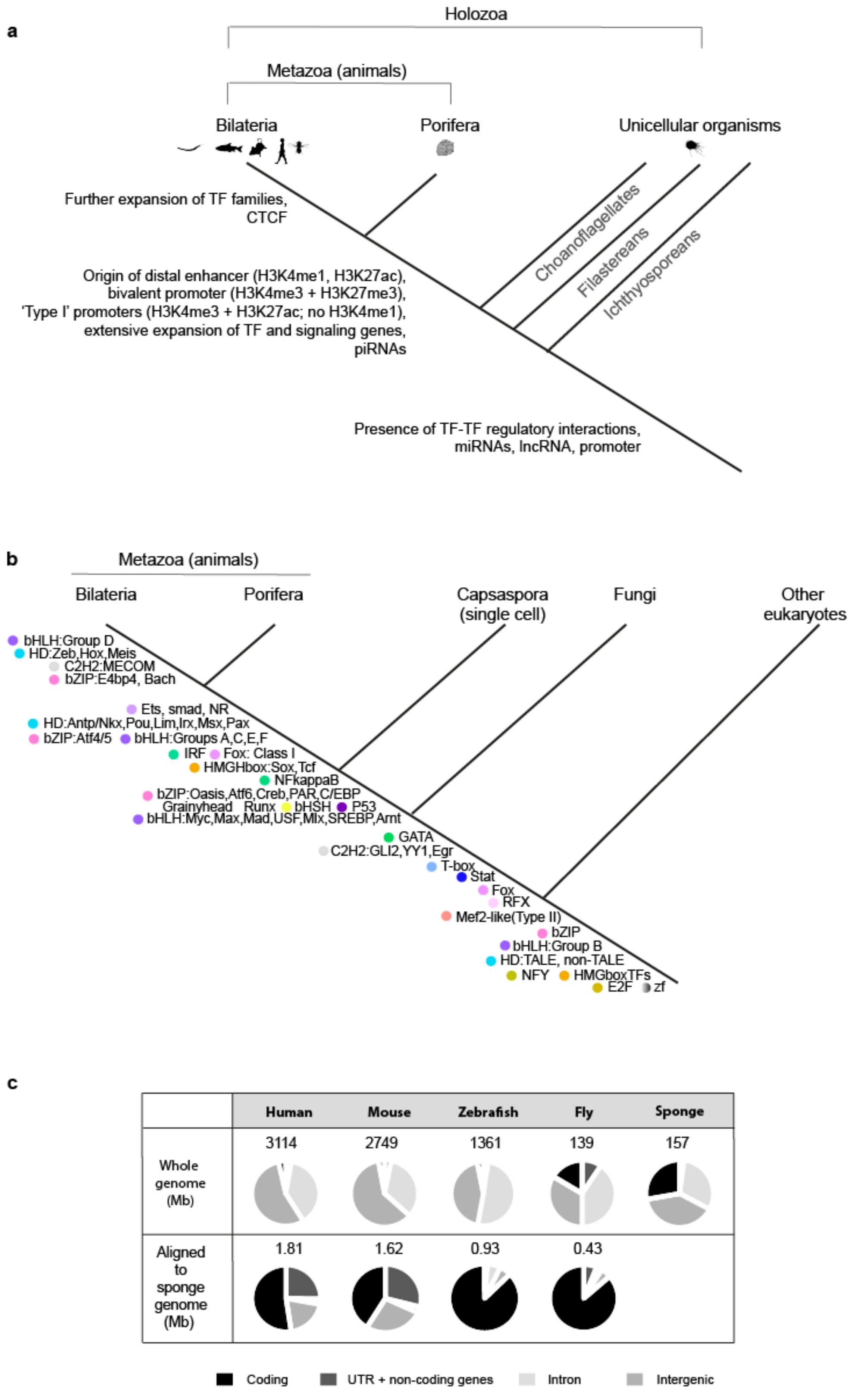

***Figure S1. Evolution of the animal regulatory genome***

**a)** Evolution of metazoan genome regulatory landscape. Based on information inferred from the genomes of extant species, distal enhancer regulation (enhancers at non-first introns and intergenic regions), and both developmental (bivalent) and differentiated cell ('Type I') promoter types appear to be animal innovations [6,32–34]. On the other hand, proximal *cis*-regulatory elements and type II promoters, which show H3K4me1, K3K4me3 and K27ac marks and label ubiquitously expressed genes, predate the metazoan lineage. **b)** The majority of TF families are conserved between Porifera and eumetazoans [5,32,35]. **(c)** First row shows animal genome sequences partitioned to different classes of sequence. The blastn algorithm was used to align the sponge genome to the genomes of other species (e-value<1e-3) and the second row shows the proportion of other genomes aligned to the sponge revealing little sequence similarity at non-protein coding regions. Repeat-masked genome sequences and full annotations for human (GRCh38.p12), mouse (GRCm38.p6/mm10), zebrafish (GRCz11) and fruitfly (BDGP6) were downloaded from Ensembl (Release 95). Locations of coding, UTR and non-coding genes, intron and intergenic distances were calculated from GFF3 files. [1]. *Amphimedon* genome assembly Aqu1 with genome annotation version Aqu2.1 was used.

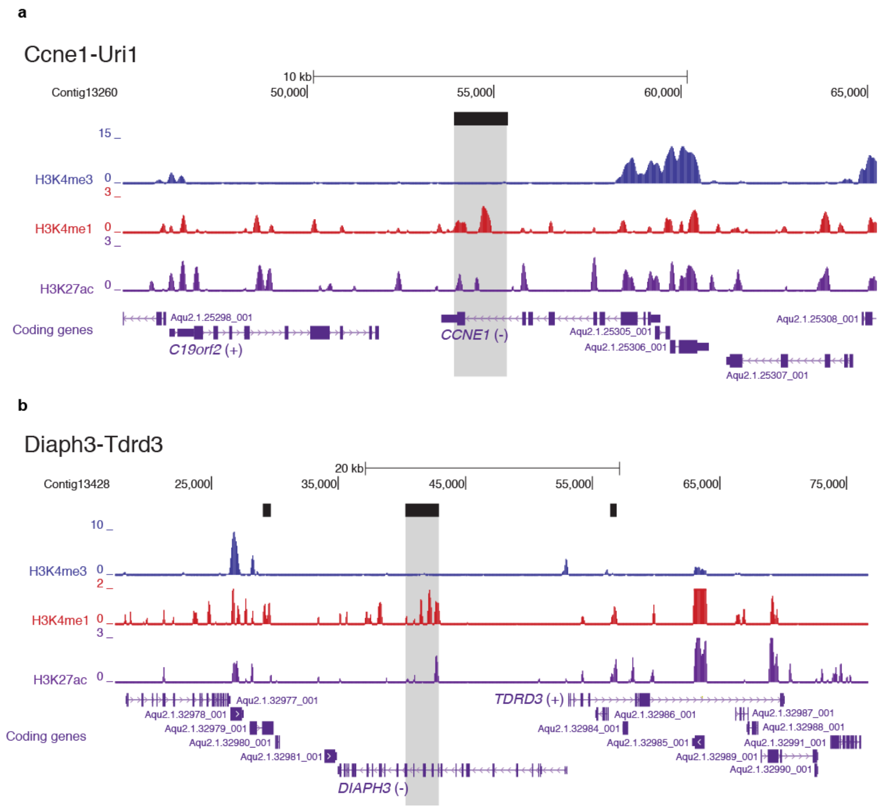

***Figure S2. Genome browser views of Ccne1–Uri and Tdrd3–Diaph3 enhancers***

Genome browser shots of **(a)** Ccne1-Uri and **(b)** Diaph3-Tdrd3. Grey region denotes where candidate enhancers are located. Tracks for H3K4me3, H3K4me1 and H3K27ac are shown [6]. H3K4me1 colocalization with H3K27ac is known to mark active enhancers in mammals.

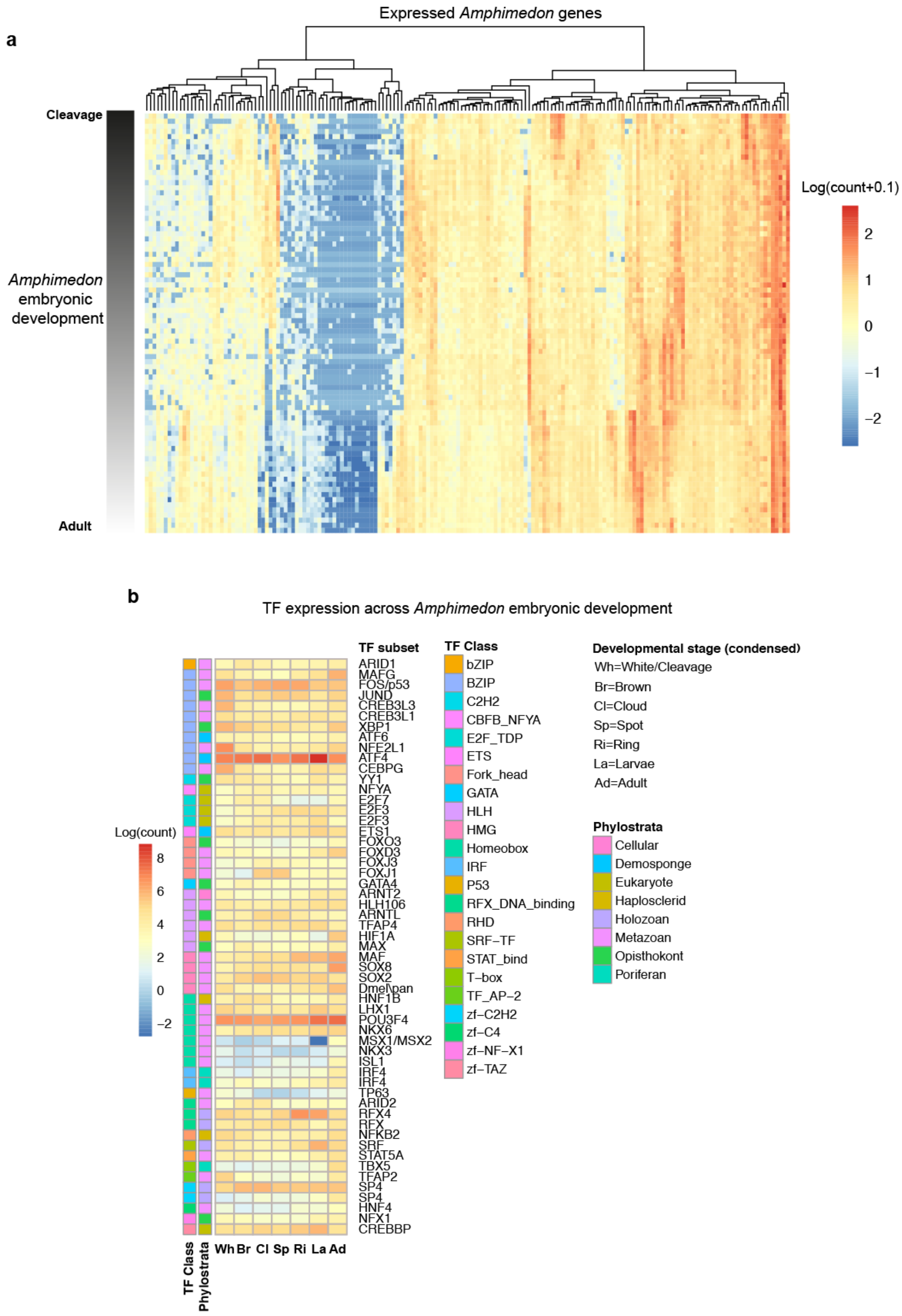

***Figure S3. Transcriptional regulation in Amphimedon queenslandica***

**(a)** A dense developmental transcriptomic time course generated using 82 CEL-seq datasets shows dynamic gene expression during development. Natural log of counts taken. Genes with a standard deviation of 0 are not shown nor used in subsequent analyses. **(b)** Sponge TFs show orthology to well-studied metazoan TFs. A subset of high confident sponge TFs are shown with information regarding their TF class, phylostrata and their expression during embryonic development. Natural log of counts taken.

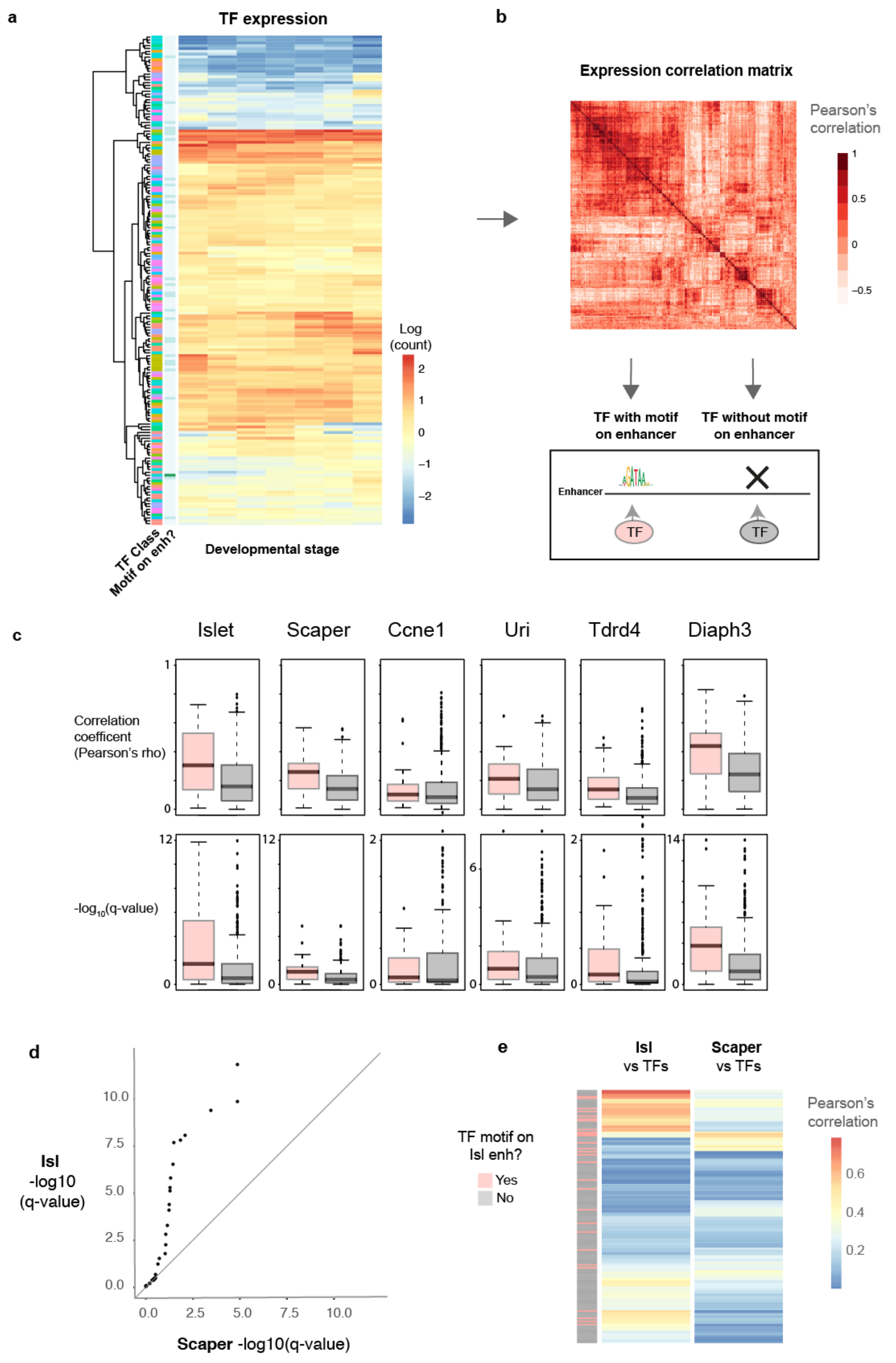

***Figure S4. Correlative relationship of TFs associated to PWM with microsyntenic genes***

**(a)** We first classified TFs based on the presence of their PWMs on an enhancer and took those expressed in at least one library in an 82 sample CEL-seq developmental transcriptomic dataset. **(b)** We then constructed a correlation matrix for all genes (Pearson's) and inferred regulatory relationships between TFs and gene targets. **(c)** Microsyntenic genes were more correlated to TFs associated to their respective enhancers than other TFs. **(d)** qqplot showing candidate sponge TFs for the Islet-Scaper enhancer, based on annotation of TF binding motifs, were better correlated with the developmental gene Islet than Scaper (n=30). q-values were obtained from Pearson's correlations of gene expression levels across sponge development from CEL-seq data [29]. **(e)** Heatmap representation of Pearson's r for the comparison described in (b) where each row represents a sponge TF.

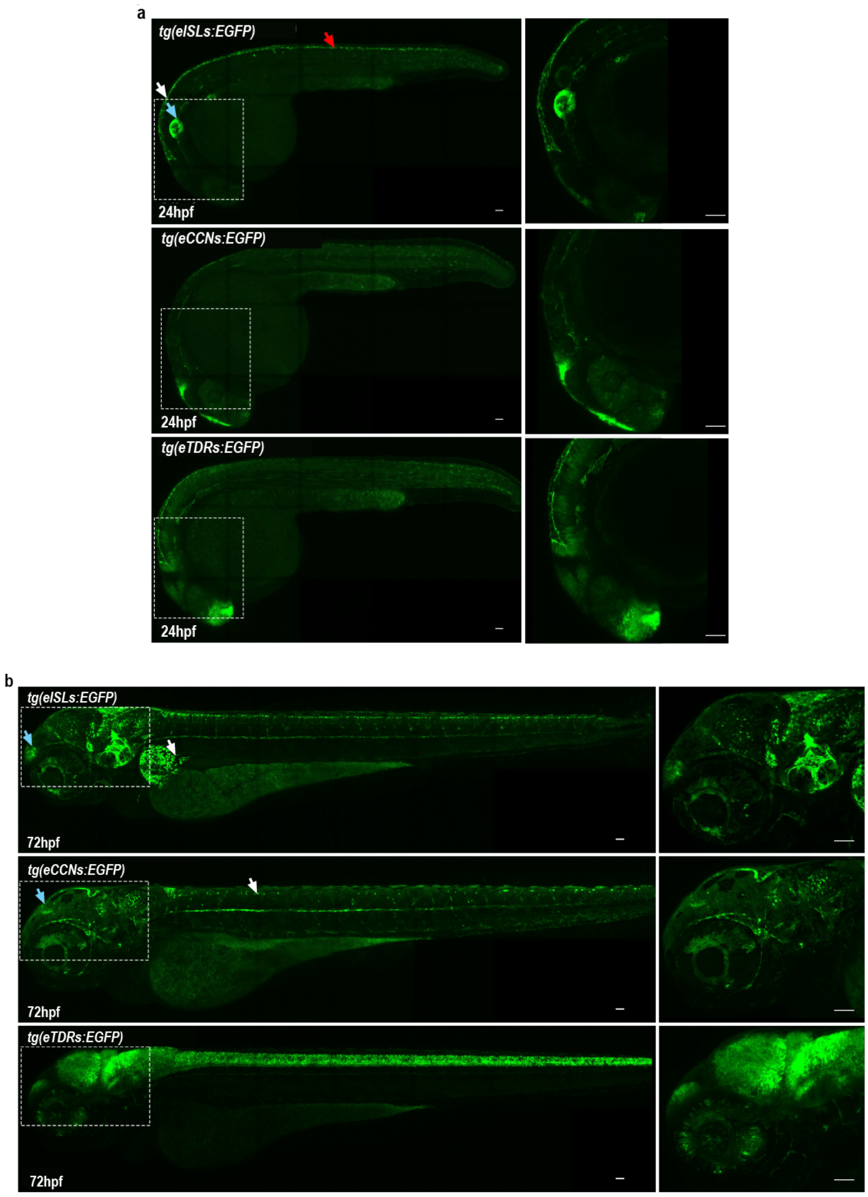

***Figure S5. Sponge enhancers show cell-type restricted activity across developmental time points in transgenic zebrafish reporter lines***

(**a**) At 24hpf, *eISLs* activity was mainly observed in the hindbrain (white arrows), otic vesicle (blue arrows) and roof plate of the neural tube (red arrows). Reporter expression in *eCCNs* was found in the epiphysis and in the telencephalon regions at 24hpf *eTDRs* drove the reporter gene expression in the forebrain, midbrain and retina at 24hpf. (**b**) At 72hpf, eISLs activity is detected in the pineal region of the forebrain (blue arrow) and in the pectoral fin (white arrow). eTDRs showed activity in the retina and in the mid-brain. Reporter expression in eCCNs show GFP signal in the mid-to-hindbrain boundary (blue arrows) and in a subset of neurones in the spinal cord (white arrows) and in facial nerve projection of motoneurons. All images shown in this figure represent the maximum intensity projection of GFP signal after confocal microscopy imaging. Scale bars: 100μM

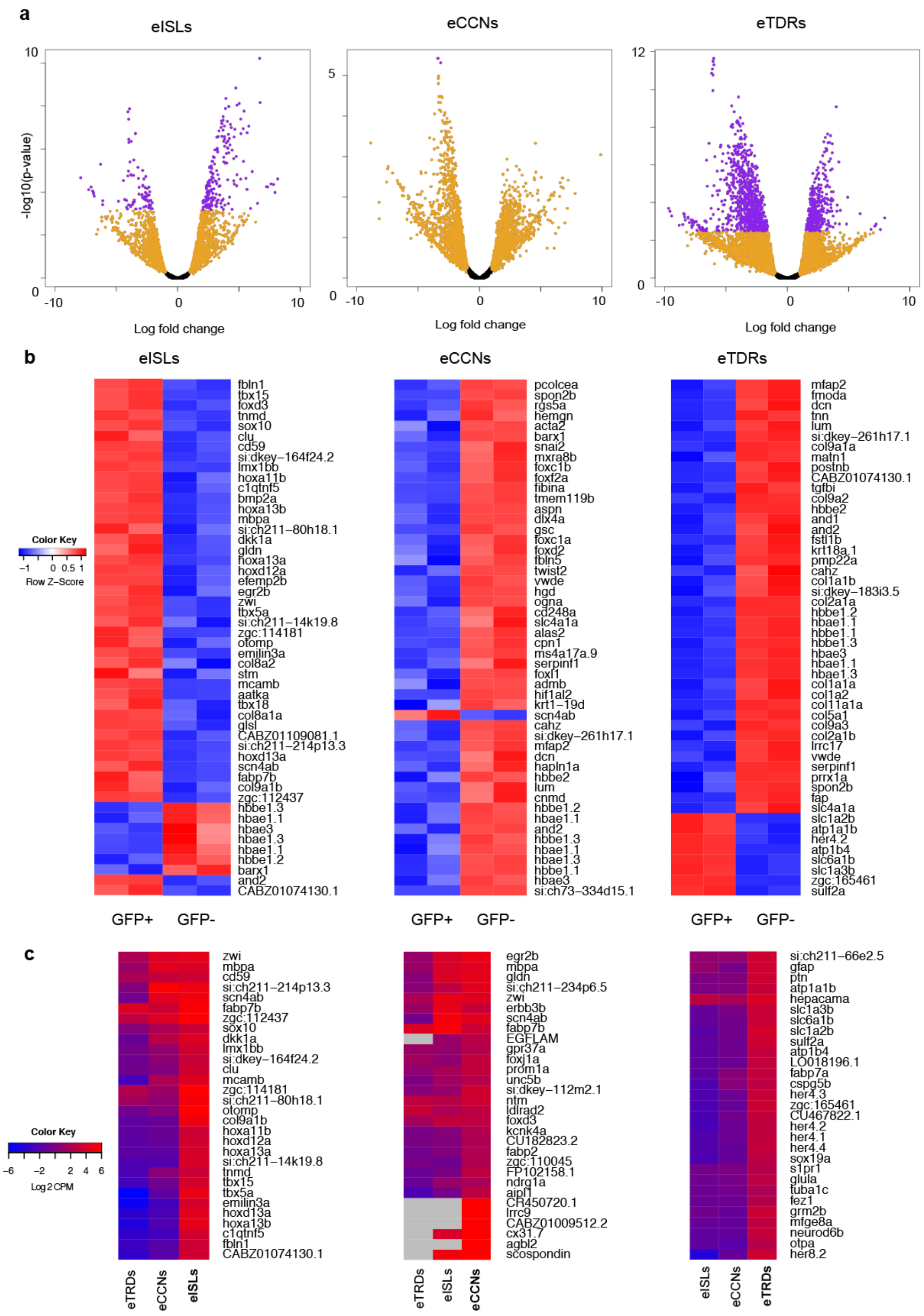

***Figure S6. Differential gene expression results between GFP positive and negative cells***

**(a)** Volcano plots shows $\log_2$ fold change on the x-axis, $-\log_{10}$(p-value) on the y-axis. Points are in orange if $\log_2$ fold change is greater than 1, and purple if $\log_2$ fold change is greater than 1 and FDR<0.05. **(b)** Heatmap shows the top 50 most differentially expressed genes between GFP positive and negative fractions for each sponge enhancer fish line. Two replicates for each condition were performed. Values show row-based z-score of $\log_2$ counts per million reads. Zebrafish gene names are shown. **(c)** Heatmaps comparing the top 30 most differentially upregulated genes for the GFP positive cells for one sponge enhancer line (in bold) relative to the other GFP positive fractions from the other sponge enhancer lines.

Values show $\log_2$ counts per million reads. Grey values denote genes that are not expressed. Zebrafish gene names are shown.

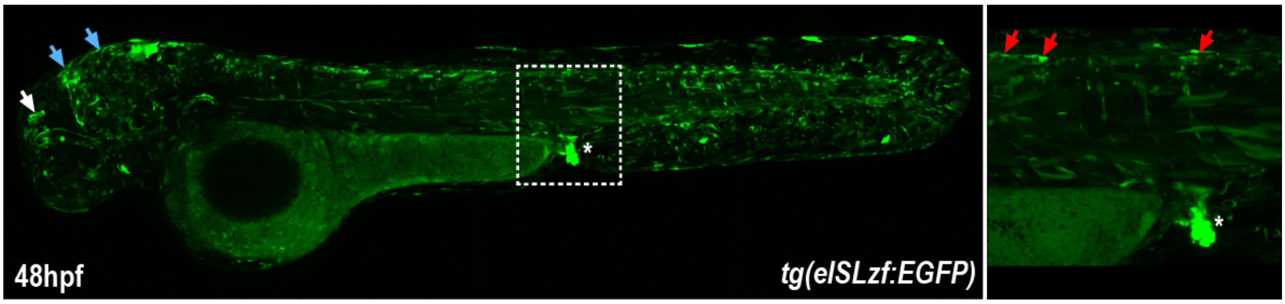

***Figure S7. Endogenous zebrafish (eISLzf:Scaper) enhancer drives cell type-specific activity in developing larvae***

Transient transgenics reporter zebrafish embryos were generated using 1-cell stage injection of a *eISLzf* putative enhancer cloned into ZED vector. This enhancer trap approach revealed activity for this regulatory element in the pineal region (white arrow), the hindbrain (blue arrows) the proctodeum (white dashed square and inset, asterisks) and some neurons of the roof plate (red arrows). The gfp activity appears mosaic at that stage since images show a F0 transient transgenic reporter fish. 11 out of 44 injected embryos displayed mosaic reporter expression The transient endogenous *eISL* lines showed a similar pattern of expression as previously reported [7]. Transgenic sequence inserted located at chr25: 32313729-32314999 (GRCz10/danRer10).

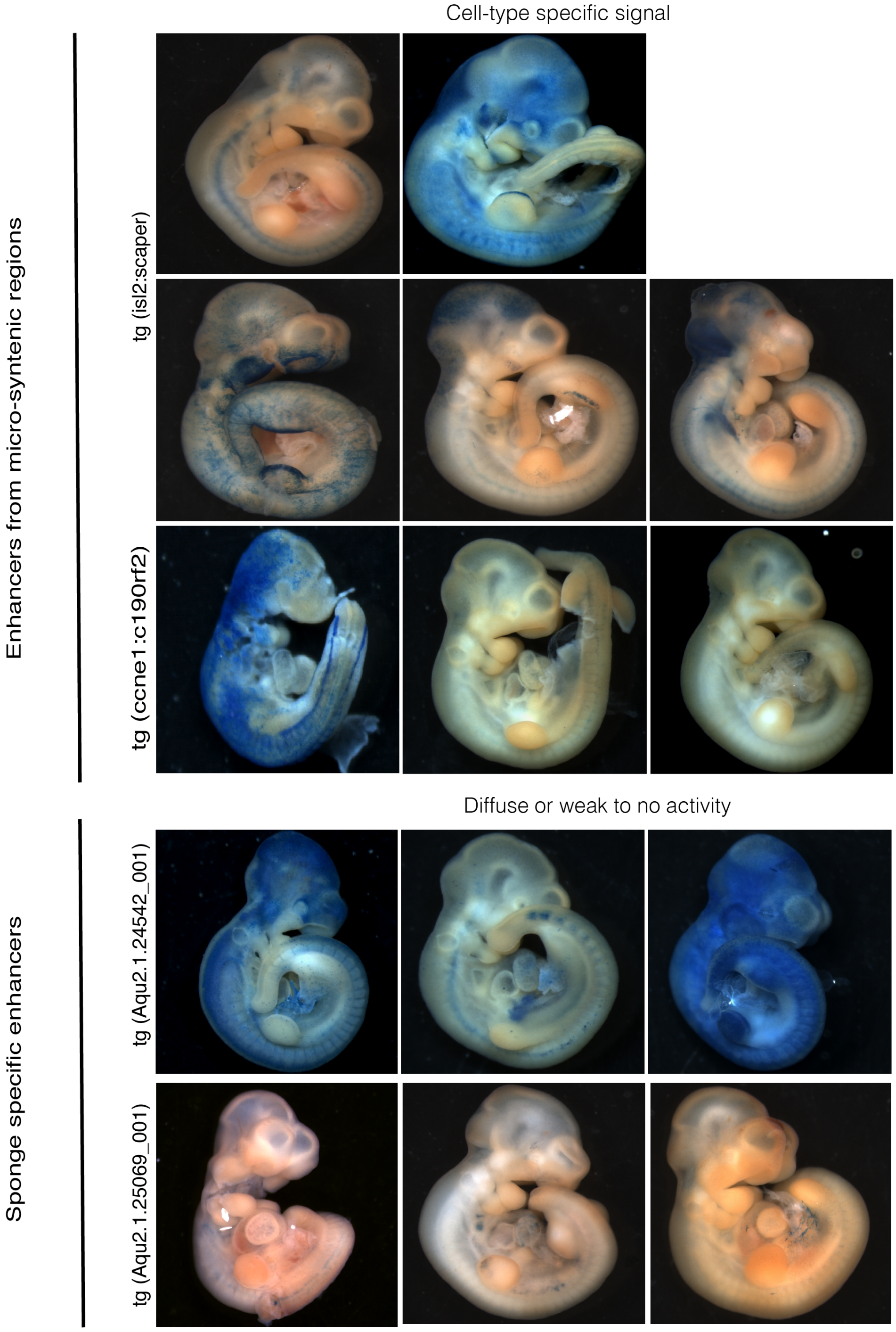

***Figure S8. Sponge enhancers from microsyntenic regions direct cell-type specific activity in mammalian embryo***

Whole mount LacZ staining of transient transgenic mouse embryos at 10.5dpc. Transient transgenic embryos were generated by pronuclei injection using sISL2:Scaper (n=6 lacZ positive out of 28) and sCcne1:C190rf2 (n=3 LacZ positive out of 25) enhancers from microsyntenic regions or Aqu2.1.25069_001 (n=3 LacZ positive out of 21) and Aqu2.1.24542_001 (n=3 Lacz positive out of 44) enhancers as sponge specific regulatory elements. While enhancer from microsyntenic region display activity in specific cell type during development, sponge-specific enhancer drive a broad or weak expression pattern of LacZ reporter gene.

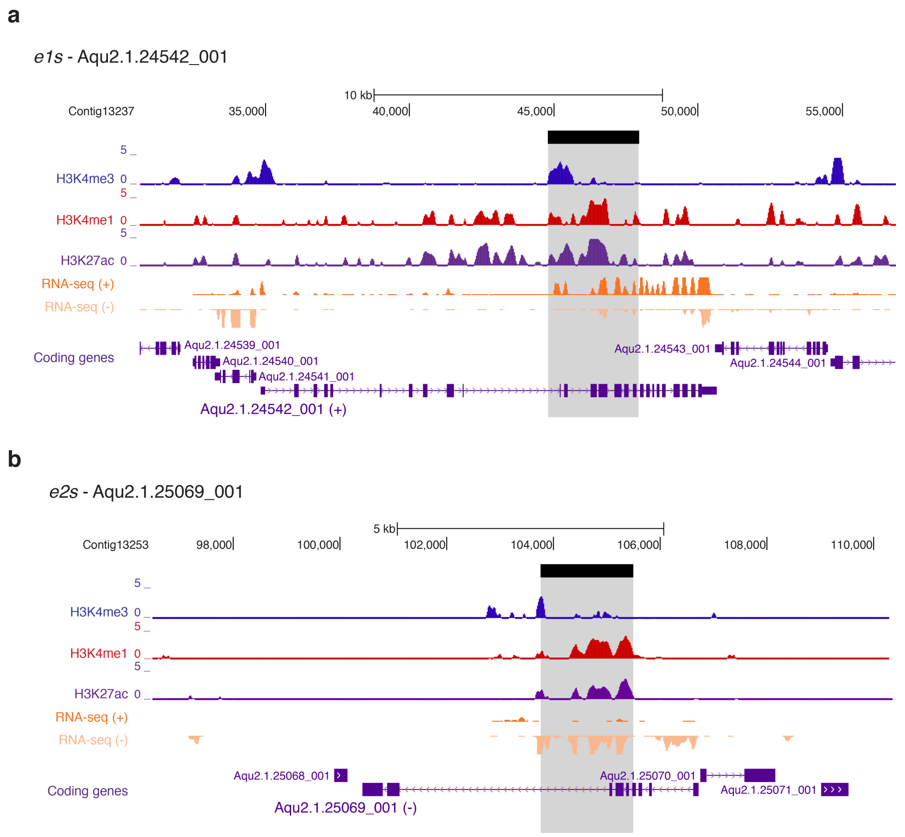

***Figure S9. Genome browser views of enhancers within sponge-specific regions***

Genome browser shots of two novel enhancers (*e1s* **(a)** and *e2s* **(b)**) located within protein-coding genes that are sponge-specific, lacking 1-to-1 vertebrate orthology. Grey regions denote where candidate enhancers are located. Tracks for H3K4me3, H3K4me1 and H3K27ac and RNA-seq expression are shown [6]. H3K4me1 colocalization with H3K27ac is known to mark active enhancers in mammals.

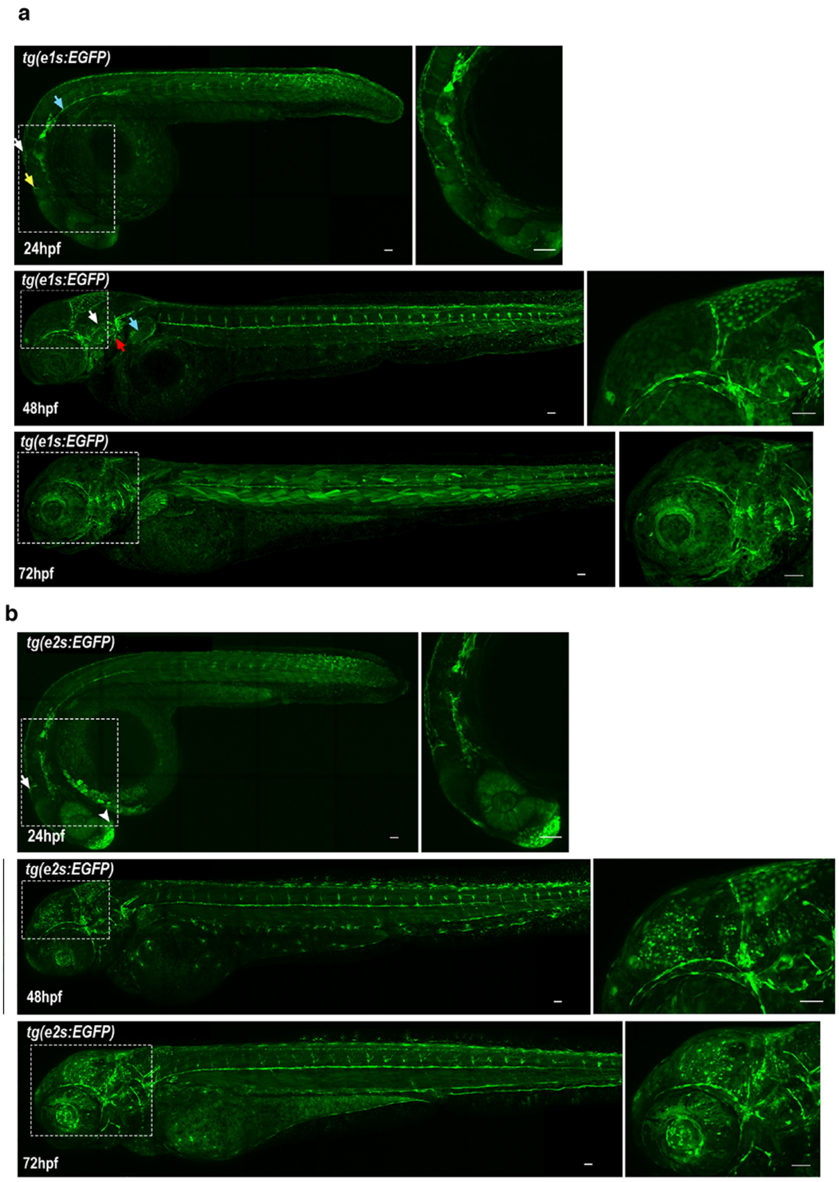

***Figure S10. Enhancers within sponge-specific regions (e1s and e2s) drove broad GFP expression pattern with variable tissue-specific activity across developmental time points***

**(a)** At 24hpf, *e1s* activity is detected in the mid-brain tectum (white arrows), the floor plate (blue arrows) and shows some weak activity in the diencephalon (yellow arrows). At 48hpf, GFP expression in the brain has mostly receded except for the neuro-epithelial cells of the hindbrain. Activity of *e1s* is observed in axon projection of facial motoneurons (red arrows), in the otic vesicle (white arrows) and the pectoral fin (blue arrow), and in Rohon-Beard neurons and caudal primary motoneurons connecting to the horizontal myoseptum. At 72hpf, *e1s* activity in the trunk has switched mostly to muscle cells whereas activity in the head remains similar to 48hpf. **(b)** At 24hpf e2s is expressed weakly in the hindbrain, the floor plate, the retina and the midbrain tectum, with most of its activity in the telencephalon (arrowheads) and diencephalon (arrows). At 48hpf, expression has switched from retina to a subset of neurons in the lens, with a broad expression pattern in the mid brain, hindbrain, the otic vesicle and the pectoral fin, In the trunk, *e2s* activity is observed in the neural tube, in caudal primary motoneurons connecting to the horizontal myoseptum and in some epithelial cell layer around the tail. At 72hpf, *e2s* activity has expanded in the eye throughout retina and lens tissues, and epithelial activity is well pronounced around the tail.

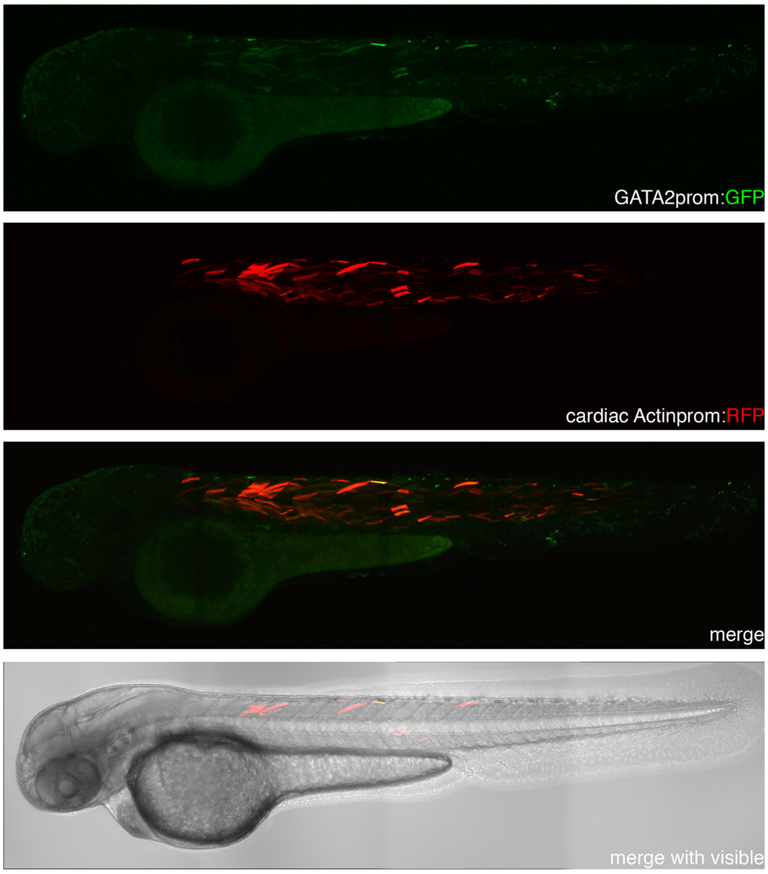

***Figure S11. Empty ZED vector display low to no background activity in transgenic fish embryos***
Transient transgenics zebrafish embryos were generated with an empty ZED vector to assess background activity of the minimal GATA2 promoter, at random integration sites, driving GFP reporter gene expression. At 48hpf, 51 out of 200 injected embryos with the empty ZED vector showed only robust RFP expression in muscle cells (in conjunction with brightfield) demonstrating successful integration and germline transmission. In a few fish, very faint GFP expression was observed only in muscle cells (GFP panel) revealing a slight background of the basal promoter activity. This signal is lost when outcrossed to an F1 generation.

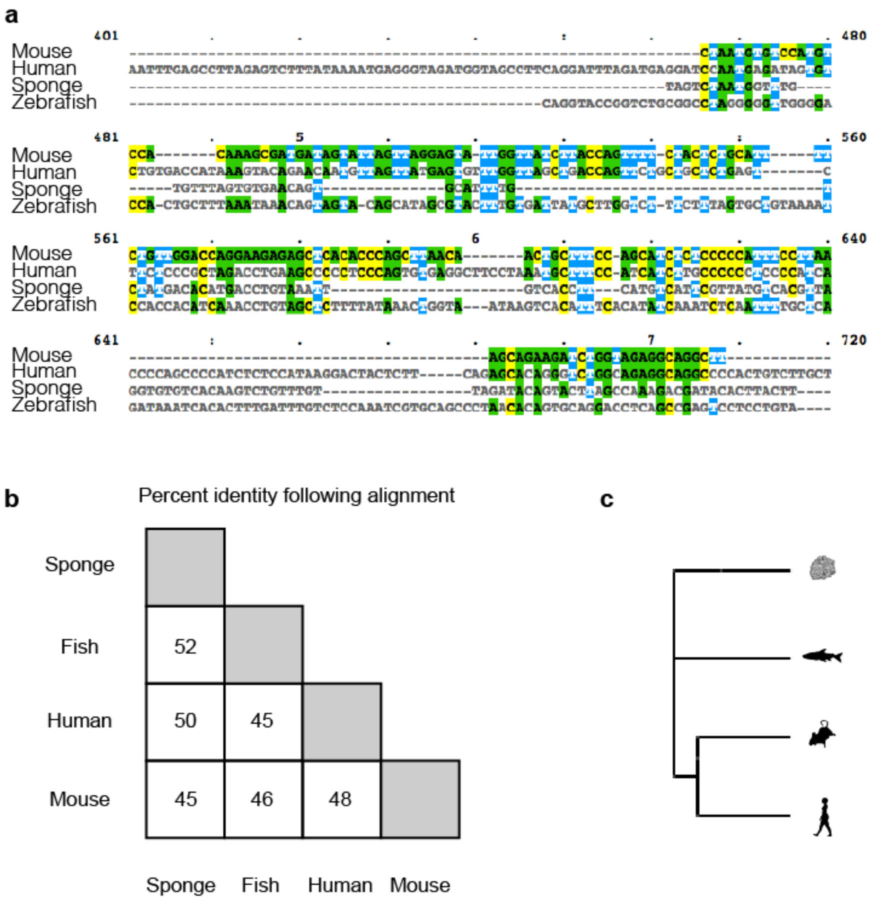

***Figure S12. Poor conservation of eISL across metazoans***

**(a)** Multiple sequence alignment of human, mouse, zebrafish and sponge candidate Islet enhancer sequences. Alignment was performed using the MUSCLE algorithm [36]. Only a subset of the full alignment is shown. This region was one of the best aligning region. **(b)** Percent sequence identity calculated following multiple sequence alignment. **(c)** Neighbour-joining phylogeny reconstructed from the alignment is unable to recapitulate correct species relationships. The aligned sequences were unable to recapitulate true phylogeny.

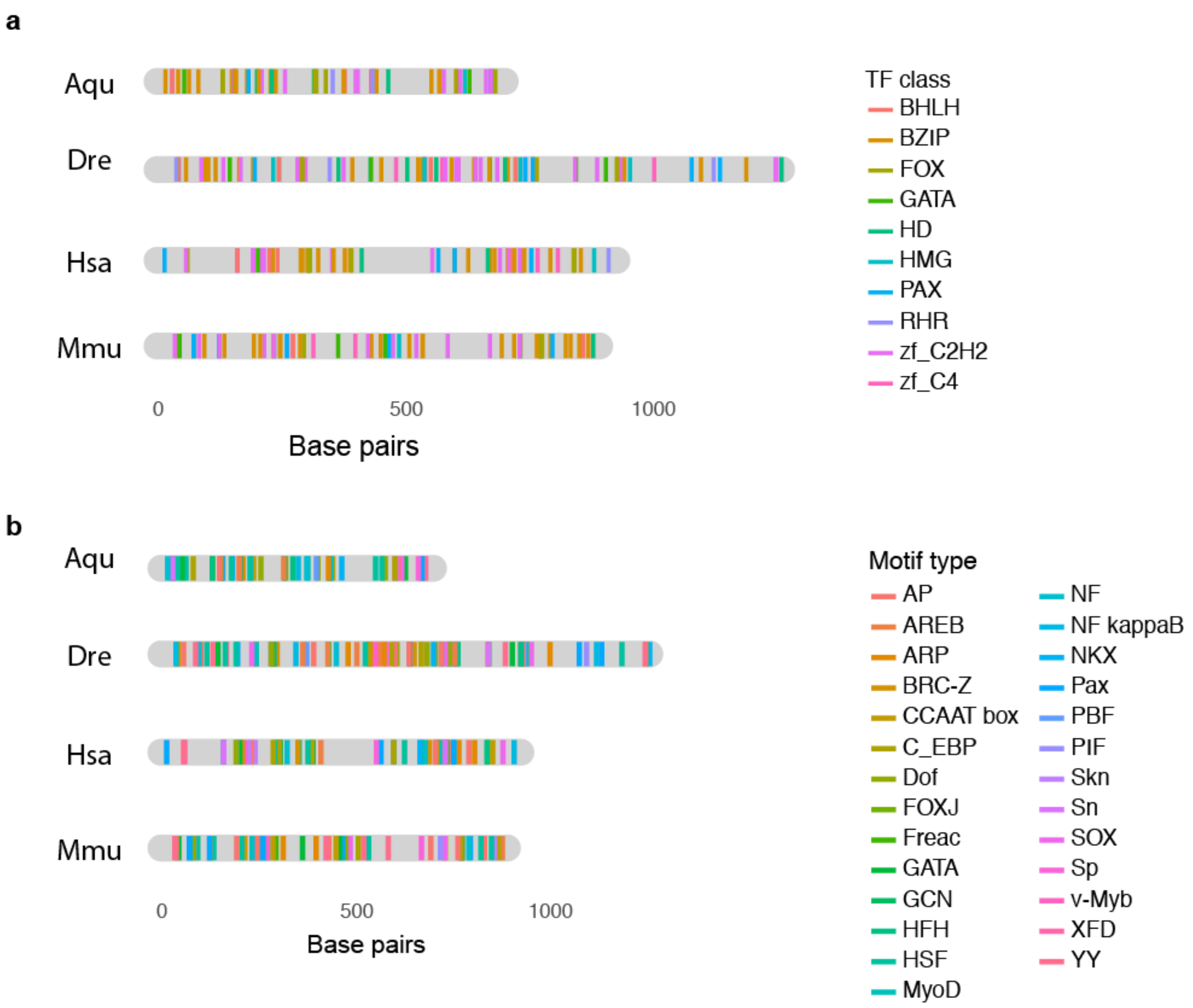

***Figure S13. Positioning of TF binding motifs along eISL sequences***

**(a)** Motif locations by the assignment of individual motifs to a broader category of TF class [37] **(b)** Motif locations by the assignment of motifs to gene families (**Methods**)

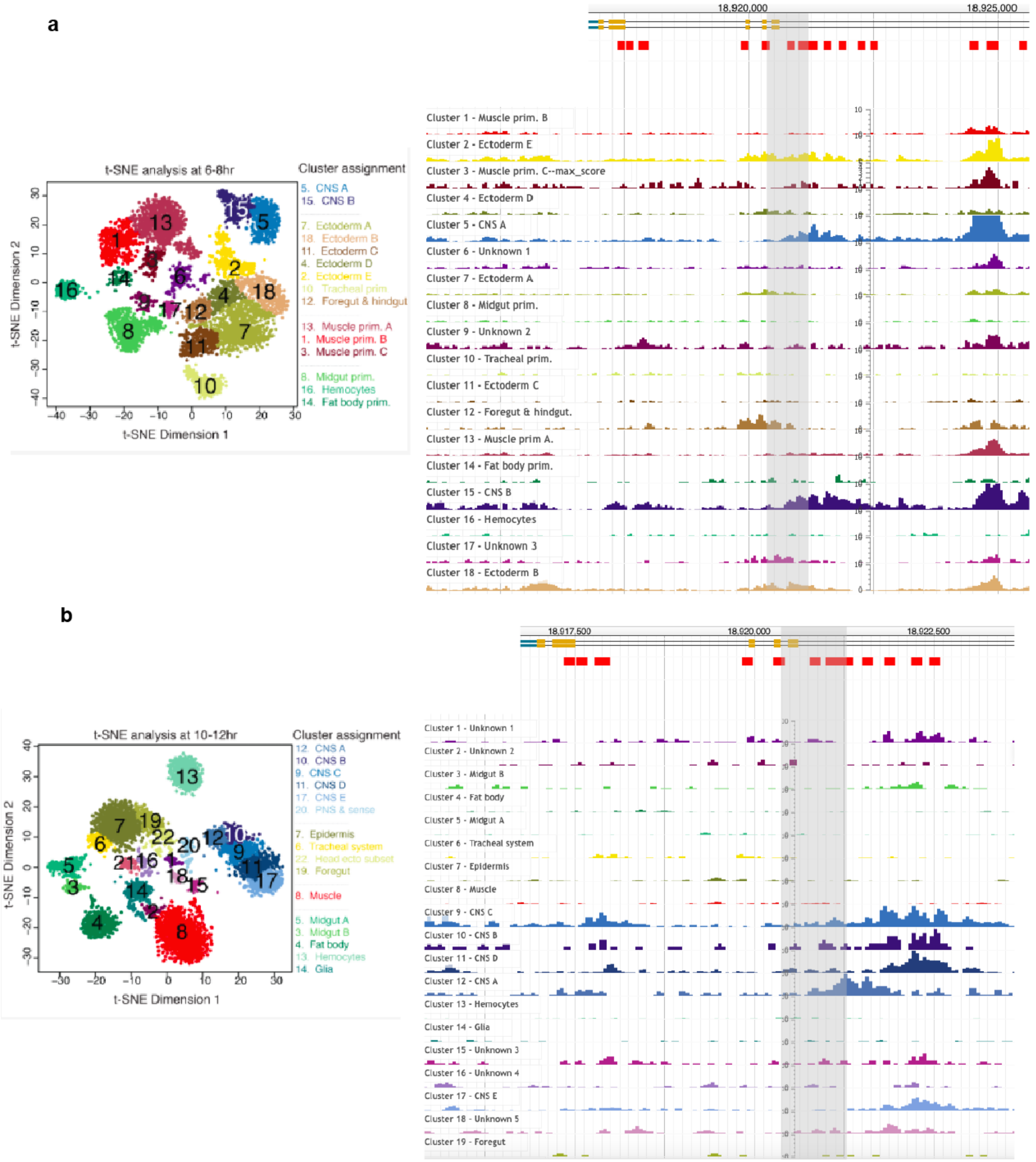

***Figure S14. Single cell ATAC-seq reveals the top fruitfly candidate ortholog to sponge eISLs is specifically chromatin accessible in neuronal cells***

We used our method to predict an ortholog to *eISLs* in *Drosophila melanogaster.* The top hit overlaps a region of open chromatin (chr2L:18920998-18921381) during fly development at 6-8 hours and 10-12 hours after egg laying. Using single cell ATAC-seq data [38], we show this region is active in **(a)** the two central nervous system (CNS) clusters (5 and 15)(FDR<4e-18) at 6-8 hr, and **(b)** at CNS cluster 12 at 6-8 hr. t-SNE plots show the cells clusters from Cusanovich et al. 2018 [38]. Genome browser tracks from http://shiny.furlonglab.embl.de/scATACseqBrowser/ .

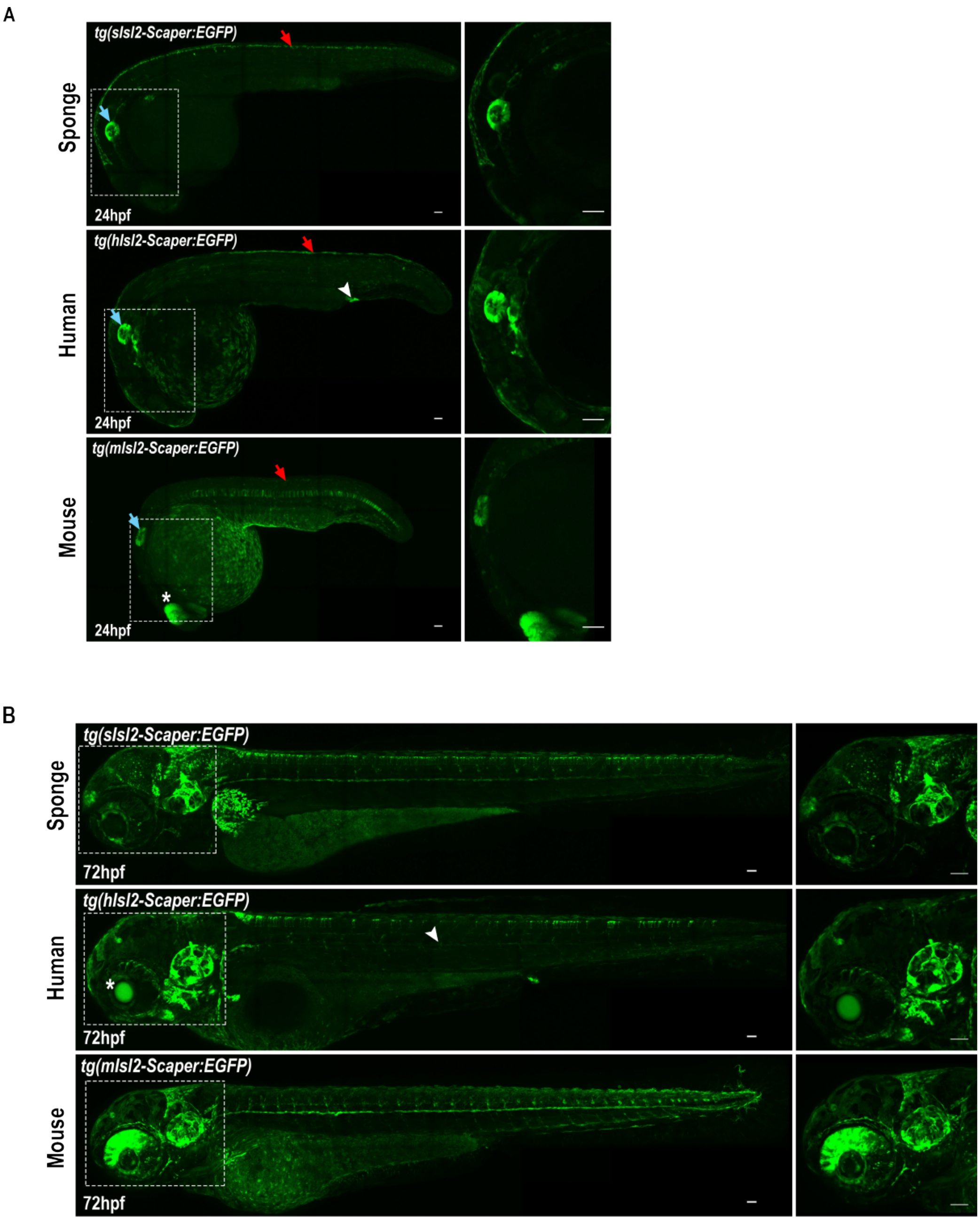

***Figure S15. Vertebrate orthologs of sponge enhancers show cell-type restricted activity across developmental time points in transgenic zebrafish reporter lines***

**(a)** Both the human and mouse orthologs (*eISLh* and *eISLm*) to the sponge Islet enhancer showed a similar but not identical activity to sponge *eISLs* in directing GFP reporter gene expression. At 24 hpf, all 3 enhancers activity is observed in the otic vesicle (blue arrows) and neurons of the roof plate (red arrows) with the mouse enhancer showing additional activity in the retina (asterisks) and the human enhancer displaying activity in the proctodeum (arrowhead). Expression in the proctodeum is similar to the expression in our zebrafish transient line (**FIGS7**) and the endogenous expression of Isl2a [7] (**b**) At 72 hpf all 3 enhancer enhancers show a consistent activity with the expression profile observed at 24hpf: otic vesicle (blue arrows) and neurons of the roof plate (red arrows). Additional activity in the in the central part of the lens (asterisks) and in endothelial cells (arrow heads) is observed for the human enhancer. By contrast, the mouse enhancer is still active in a polarised manner in a sub-population of retinal neurons.

a

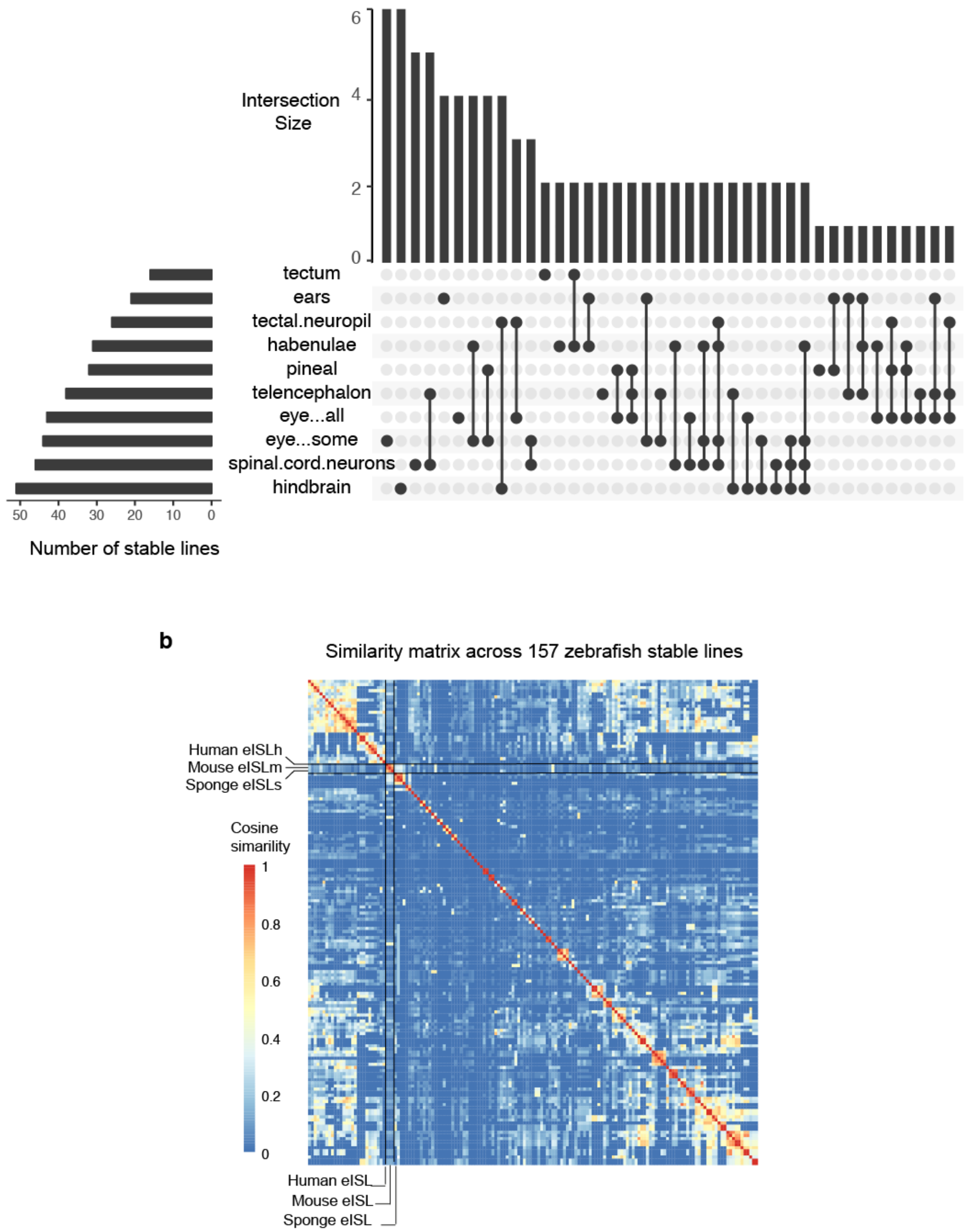

***Figure S16. Quantitative anatomical comparison across enhancer trap lines supports strong similarity between Isl enhancers in human, mouse and sponge***

**(a)** The most frequent combinations of anatomical regions where GFP reporter expression was detected based on 154 zebrafish lines from enhancer trap assay [9]. Although the putative Islet enhancers showed activity in some common tissue types in the assay (e.g. hindbrain, eye) (**FIGS18**), the combination of expression patterns are highly unique as shown in (b). **(b)** A similarity matrix derived from GFP coexpression patterns across 157 zebrafish lines, including the human, mouse and sponge ISL enhancers. The ISL lines are labeled and showed a much higher similarity to each other than other assayed enhancers. To maximize consistency, imaging of the fish was performed under similar settings and at a similar developmental time (3 dpf).

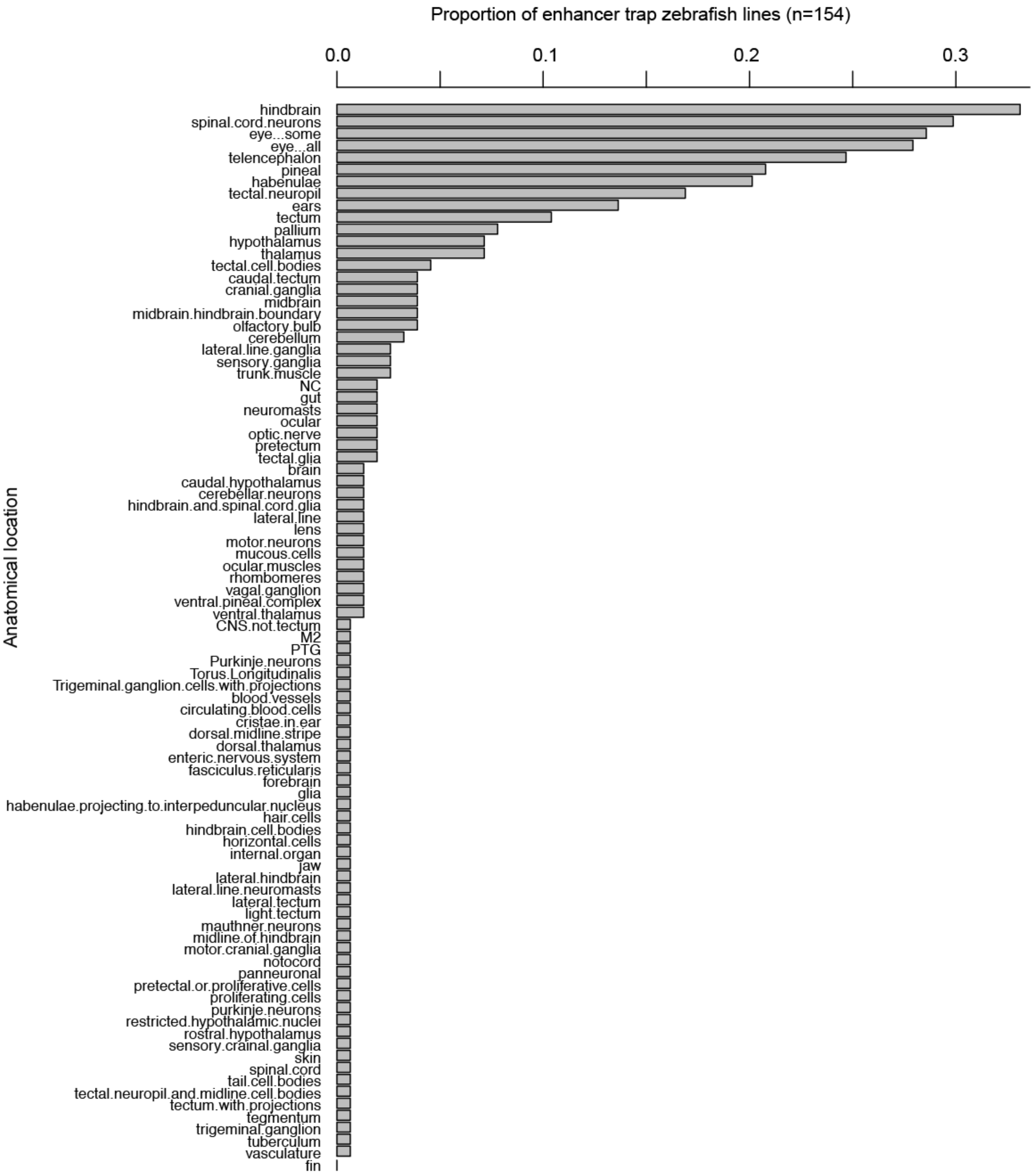

***Figure S17. Histogram of 88 anatomical zebrafish locations showing GFP reporter activity in the published 154 zebrafish stable lines from zebrafish enhancers.***

Processed data showing the frequency of enhancer activity across zebrafish tissue of 154 zebrafish enhancers in stable lines (**Methods**)[9].
